# Genetic architecture of gene regulation in Indonesian populations identifies QTLs associated with local ancestry and archaic introgression

**DOI:** 10.1101/2020.09.25.313726

**Authors:** Heini M. Natri, Georgi Hudjashov, Guy Jacobs, Pradiptajati Kusuma, Lauri Saag, Chelzie Crenna Darusallam, Mait Metspalu, Herawati Sudoyo, Murray P. Cox, Irene Gallego Romero, Nicholas E. Banovich

**Affiliations:** Center for Evolution and Medicine, School of Life Sciences, Arizona State University, Tempe 85281, AZ, USA; The Translational Genomics Research Institute, Phoenix 85004, AZ, USA; Statistics and Bioinformatics Group, School of Fundamental Sciences, Massey University, Palmerston North 4410, New Zealand; Centre for Genomics, Evolution and Medicine, Institute of Genomics, University of Tartu, Tartu, 51010, Estonia; Leverhulme Centre for Human Evolutionary Studies, Department of Archaeology, University of Cambridge, Cambridge, CB2 1QH, United Kingdom; Complexity Institute, Nanyang Technological University, Singapore, 637460; Laboratory of Genome Diversity and Disease, Eijkman Institute for Molecular Biology, Jakarta 10430, Indonesia; Institute of Genomics, University of Tartu, Tartu, 51010, Estonia; Melbourne Integrative Genomics, University of Melbourne, Parkville 3010, Australia; School of BioSciences, University of Melbourne, Parkville 3010, Australia; Centre for Stem Cell Systems, University of Melbourne, Parkville 3010, Australia

**Author notes:** **Corresponding author information**, Murray P. Cox, Irene Gallego Romero, Nicholas E. Banovich.

**Keywords:** Genetic variation, Gene regulation, eQTL, QTL, Neandertal, Denisovan, Indonesia

## Abstract

Lack of diversity in human genomics limits our understanding of the genetic underpinnings of complex traits, hinders precision medicine, and contributes to health disparities. To map genetic effects on gene regulation in the underrepresented Indonesian population, we have integrated genotype, gene expression, and CpG methylation data from 115 participants across three island populations that capture the major sources of genomic diversity on the region. In a comparison with a European dataset, we identify 166 uniquely Indonesia-specific eQTLs, highlighting the benefits of performing association studies on non-European populations. By combining local ancestry and archaic introgression inference eQTLs and methylQTLs, we identify regulatory loci driven by modern Papuan ancestry as well as introgressed Denisovan and Neanderthal variation. GWAS colocalization connects QTLs detected here to hematological traits. Our findings illustrate how local ancestry and archaic introgression drive variation in gene regulation across genetically distinct and in admixed populations.

## Introduction

As we move into the age of precision medicine, the systematic under sampling of global genetic diversity limits our ability to broadly apply biomedical research efforts across diverse ethnicities and population backgrounds (Duncan et al. 2019; Landry et al. 2018). Indeed, the vast majority of human genomics studies to date have been conducted in individuals with European ancestry, who account for a minority of the global population (Sirugo et al. 2019). To gain a comprehensive understanding of the genetic architecture of complex disease, it is critical to expand human genomics studies into diverse populations. Collection of multi-modal genomic data from traditionally undersampled populations will also allow for the mapping of genetic associations with molecular phenotypes and integration with genome-wide association studies (GWAS).

The Indonesian archipelago is one such undersampled region. Genetically and geographically structured, with a genomic cline of Asian to Papuan ancestry stretching from west to east, it is the fourth largest country in the world by population, home to 267 million people. To investigate the effects of modern and archaic local ancestry on gene regulation in Indonesians, we present the first maps of expression QTLs (eQTLs) and DNA methylation QTLs (methylQTLs) in 115 Indonesian individuals drawn from three island populations that capture the major genomic axes of diversity across the region.

## Results

### Patterns of modern local ancestry and archaic introgression vary across the three study populations

To contextualize the genetic diversity of the three study populations, we clustered the 115 Indonesian samples (Figure 1a) using principal component analysis (PCA) of genotype data, along with European and Han Chinese samples from the 1000 Genomes project. The first two principal components clearly separate the three study populations (Figure 1b). The Mentawai, representative of East Asian ancestry, cluster closest to mainland Chinese populations, whereas the Korowai, representative of Papuan ancestry, cluster distinctly from all other populations. Individuals from Sumba — a near equal mixture of the two ancestries — cluster between Mentawai and Korowai, as expected (Natri et al. 2020).

**Figure 1.**
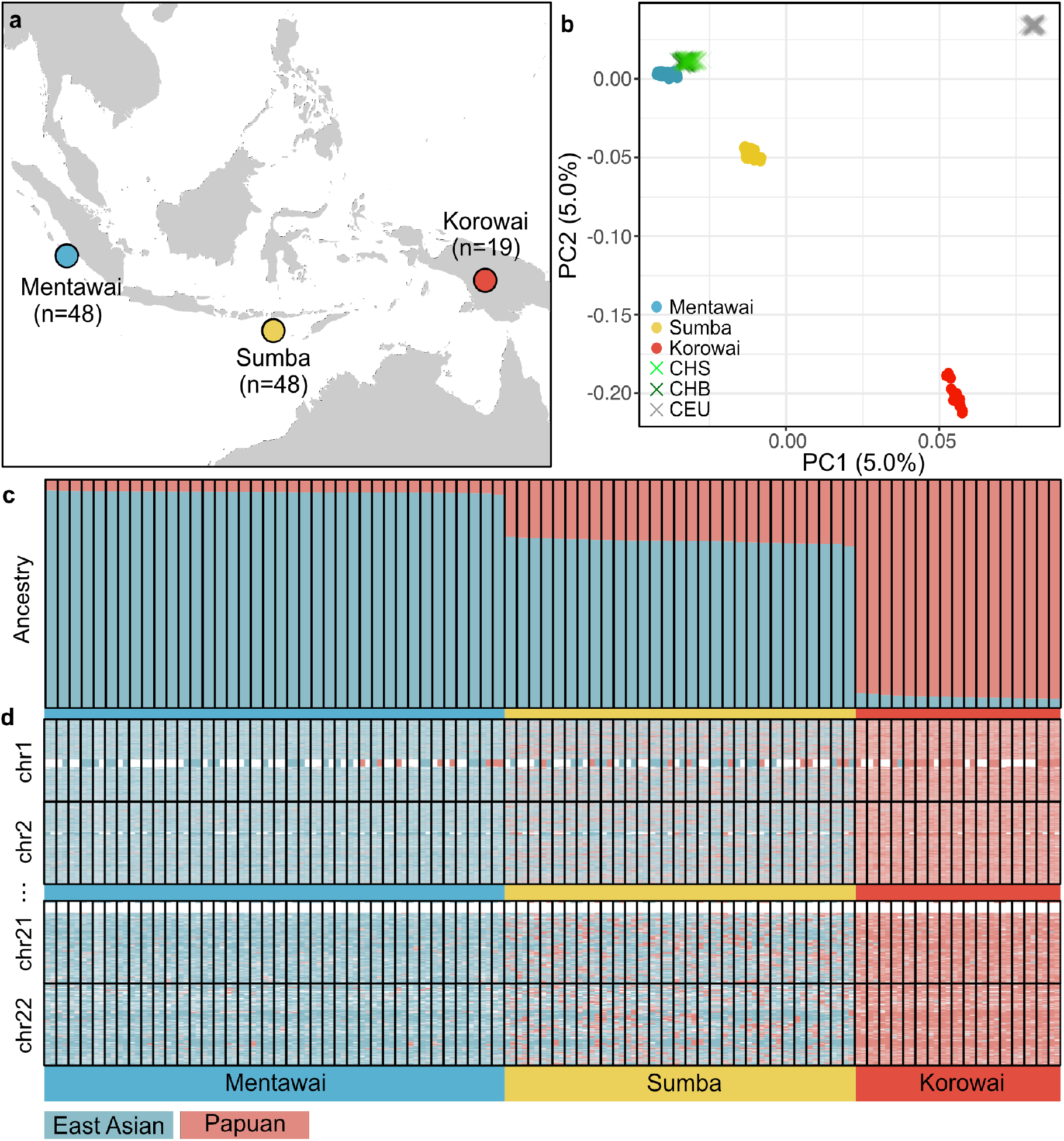
Patterns of genetic ancestry across three Indonesian island populations. **a:** Map of the sampling locations of the three study populations: Mentawai, blue; Sumba, yellow; Korowai, red. Numbers of samples used in the QTL analyses are indicated. **b:** Principal Components calculated from the genotype data of the three study populations, as well as Han Chinese from Beijing (CHB), Southern Han Chinese (CHS), and individuals with Northern and Western European ancestry (CEU) from the 1000 genomes project. **c**, **d**: Estimates of global (**c**) and local (**d**) ancestry across 73 individuals (bars) with available WGS data. **c** shows proportions (y-axis) of East Asian and Papuan ancestry across all chromosomes. **d** shows patterns of local ancestry across the two haplotypes in each individual across chromosomes.

We genotyped our samples using two separate platforms, whole-genome sequencing (WGS, n = 73) and a genotyping array (n = 42; Methods, Supplementary Table 1). Using only the complete genome sequences, we inferred patterns of global and local ancestry (LA) and archaic introgression across the three populations. On average, the proportion of the genome for which we can make a confident ancestry assignment is 80% for Mentawai, 71% for Sumba, and 85% for Korowai. The average proportion of ancestry-called individual haploid genomes assigned as Papuan is 5.3% in Mentawai, 26.8% in Sumba, and 95.0% in Korowai (Figure 1c,d). In addition, we were able to identify Denisovan-introgressed haplotypes covering, on average, 0.13%, 0.48%, and 1.44%, of each haploid genome in Mentawai, Sumba, and Korowai, respectively, consistent with previous studies showing a high frequency of Denisovan sequence in Korowai (Jacobs et al. 2019). Proportions of inferred Papuan ancestry and Denisovan introgression are highly correlated (Pearson’s r = 0.995, Extended Data Figure S1). Further, we identified Neanderthal-introgressed haplotypes covering on average 1.08%, 1.19%, and 1.40% of each haploid genomes from the three study populations.

### Genetic determinants of gene expression and CpG methylation levels in Indonesia

We used a linear regression-based approach to identify genetic variants associated with changes in expression (eQTL) and methylation (methylQTL) levels (Methods). At an FDR of 0.01, we detect a total of 1,975 significant *cis*-eQTLs (Supplementary File 1) and 48,014 *cis*-methylQTLs (Supplementary File 2). The majority of eQTLs and methylQTLs are located in non-coding parts of the genome (Methods, Figure 2a,b), and are enriched among transcriptionally active histone marks and accessible chromatin, and mostly depleted on marks associated with heterochromatin and repression of transcription across three blood cell lines (Figure 2c,d, Extended Data Figure 2).

**Figure 2:**
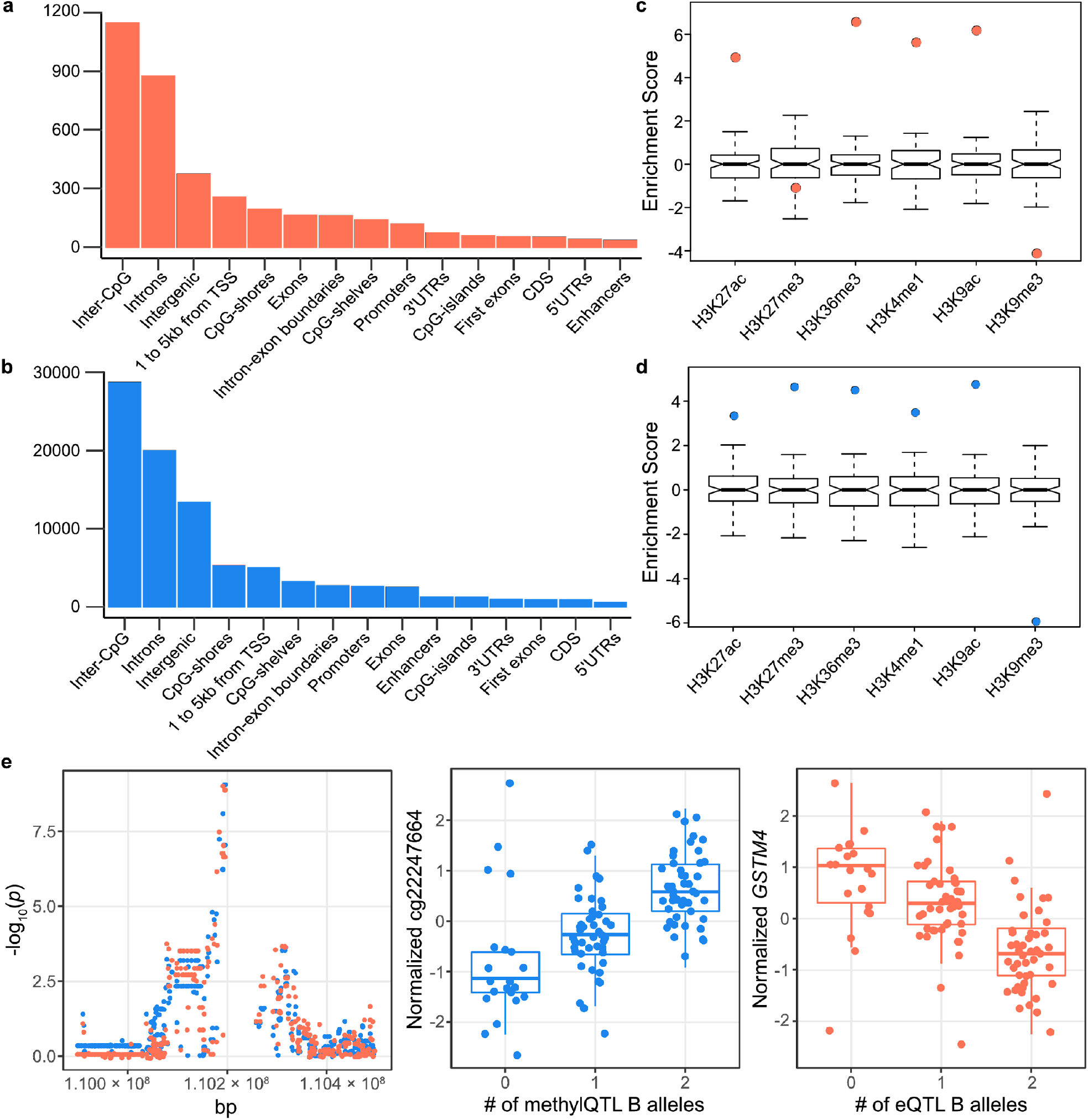
Qualities of QTLs called across 115 individuals. **a**, **b**: Genomic annotations of eQTLs (**a**) and methylQTLs (**b**). **c**, **d**: Enrichment of eQTLs (**c**) and methylQTLs (**d**) among histone marks derived from primary mononuclear cells from peripheral blood in the Epigenomics Roadmap project. Enrichment was tested against a background null-distribution of 100 sets of variants. **e**: An example of a colocalized eQTL-methylQTL pair exhibiting an opposing effect direction on the target trait. The left-side plot shows the −log_10_(*p*) of the associations between variants in *cis* and *GSTM4* expression (orange) orcg22247664 methylation (blue). The middle plot shows the relationship between the top-SNP genotype and cg22247664 methylation, and the right-side plot the relationship between the top-SNP genotype and *GSTM4* expression.

We integrated the eQTL and methylQTL calls to gain insight into how genetic regulation of CpG methylation contributes to the regulation of gene expression (Supplementary Note 1). We tested for colocalization between 4,639 pairs of CpGs and genes using a Bayesian method (Methods, Supplementary Note 1). Over a wide range of prior probabilities, 720 (15.5%) of the tested pairs show robust support for a common causal variant (Methods, Supplementary Figure S1, Supplementary File 5), corresponding to 621 unique CpGs and 222 unique genes. We had previously identified 80 (36.0%) of the 222 genes as showing a negative correlation between expression and promoter methylation levels (Natri et al. 2020). CpGs located on promoters are more likely to show an opposite direction of effect with the gene than CpGs located outside regulatory regions (Fisher’s test *p* = 3.835×10^−6^, Figure 2e, Supplementary Note 1, Supplementary Figure S2). These findings illustrate the relationship between genetically regulated promoter methylation and gene expression, highlighting the benefits of integrating multiple types of molecular data for a better understanding of the gene regulatory machinery.

### Sharing of eQTLs between Indonesian and European populations

The majority of eQTL studies have been carried out in European populations and do not capture global genetic diversity. To better understand the impact of ancestry on the genetic architecture of gene regulation, we compared eQTLs detected here (at a relaxed FDR of *p*< 0.10 to account for differences in power) with those identified in a comparable European dataset (Lepik et al. 2017). Of the 3,049 genes tested for colocalization (Methods), 48.9% (1,489) had some evidence of colocalization and 17.9% (547) exhibited robust evidence of colocalization with a wide range of prior probabilities, suggesting a high probability of a single common causal variant shared across the populations (Figure 3a,d). Concordant with previous reports, eQTLs that are shared between populations exhibit larger effect sizes than other eQTLs (t-test *p* = 9.4×10^−12^), and most (87%) shared eQTLs show the same direction of effect in both populations (Stranger et al. 2012) (Figure 3a, Extended Data Figure 3, Supplementary Note 2). Conversely, 50.3% of genes (1,660) exhibited no evidence of co-localization even with relaxed thresholds (Methods). However, 1,494 were associated (nominal *p* < 5×10^−4^) with an eQTL in Europeans. Out of these, 440 also showed nominally significant *p*-values for the Indonesian top-SNPs. 512 genes did not show evidence for colocalization and had the same top-SNPs testable in both populations, indicating that these genes are regulated in *cis* by different causal variants in each population. The absolute effect sizes for the Indonesian top-SNPs of these 512 genes differ between populations (paired t-test *p* < 2.2×10^−16^), indicative of fundamentally population specific effects (Figure 3b). Furthermore, the minor allele frequencies (MAF) of the Indonesian top-SNPs were significantly different between populations (paired t-test *p* < 2.2×10^−16^, Figure 3c), likely resulting in reduced power to detect these associations in the European data. These finding illustrate that analyses on diverse populations can lead to the discovery of new potentially causal associations even with moderate sample sizes. Despite the smaller sample size and reduced power to detect small effect sizes in our data, we identify 166 Indonesian-specific eGenes that do not show any evidence of colocalization and were not nominally associated with variants in *cis* in the European population (Figure 3a,e, Supplementary Note 2). Several of these genes are clinically relevant, including cancer-related *NDC80* (Figure 3e), *NRAS, BRMS1*, and *MRC2* (Supplementary Table 4). When compared to a background set of all tested genes, Indonesia-specific eGenes are enriched among canonical pathways involved in apelin signaling, platelet activation, Rap1 signaling, and melanogenesis (Supplementary Table 5). These results highlight the importance of studies on non-European populations for a better understanding of genetic regulation of molecular traits.

**Figure 3.**
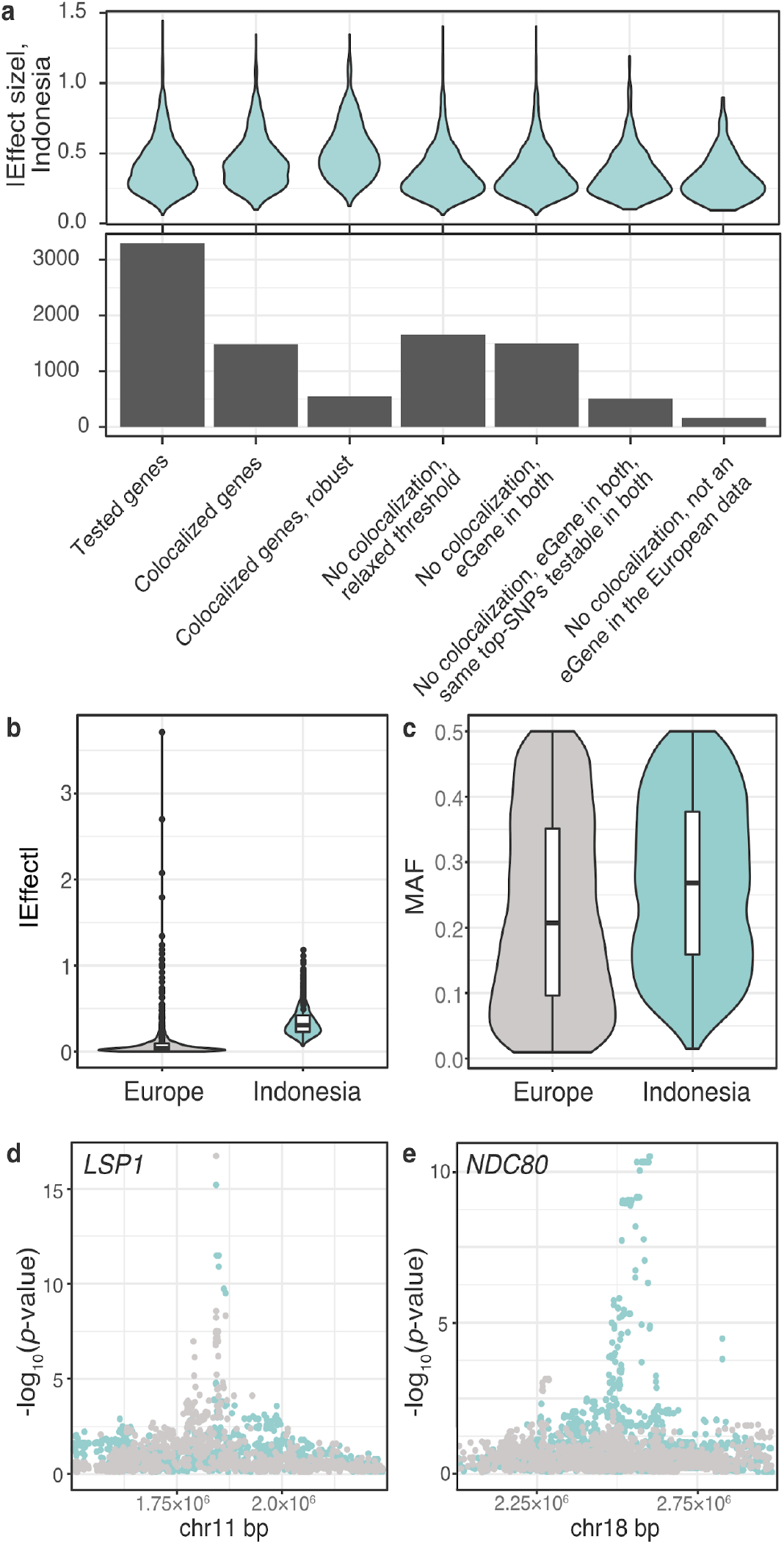
Sharing of eQTLs between Indonesian and European populations. **a:** A subset of Indonesian eQTLs shows evidence of colocalization with European eQTLs. Violin plots of absolute effect sizes for the top-SNPs in the Indonesian data for genes belonging to each group. The barplot shows the numbers of genes belonging to each group. **b, c:** The Indonesian top-SNPs for genes that do not show evidence of colocalization exhibit differences in absolute effect sizes (**b**, paired t-test *p* < 2.2×10^−16^) and MAFs (**c**, paired t-test *p* < 2.2×10^−16^) between populations. **d**, **e**: Examples of a colocalized gene, *LSP1* (**d**), and a gene that has an eQTL in the Indonesian data, but not in the European data, *NDC80* (**e**). Manhattan plots show the chromosomal locations of tested SNPs on the x-axis and the −log_10_(*p*-values) of each SNP on the y-axis. Indonesian = turquoise, European = gray.

### Subsets of methylQTLs and eQTLs are largely driven by modern local ancestry and archaic introgression

In addition to differences between Indonesians and Europeans, we sought to understand the extent to which the two distinct sources of local ancestry (LA) in modern Indonesians, as well as introgression from archaic hominins, have impacted gene regulatory architecture. We examined the haplotype background on which our observed QTLs occur and asked whether there was a relationship between the inferred ancestral source of the genotype and expression/methylation levels (Methods, Figure 4a,d,e). We find 9, 2, and 31 instances where variance in eQTL genotype is largely driven (*R^2^* > 70%) by modern LA, archaic Denisovan introgression, and archaic Neanderthal introgression, respectively (Figure 4b), directly linking ancestral alleles to gene expression differences between individuals. Similarly, we find 301, 112, and 477 instances where the methylQTL genotype is driven by modern LA, Denisovan introgression, and Neanderthal introgression (Figure 4c). In total, 2.1% of eQTLs and 2.29% of methylQTLs are driven by modern LA or archaic introgression. Of the 9 and 373 unique Papuan-driven QTL target genes and CpGs, 7 (77.8%) and 270 (72.4%) were previously identified (Natri et al. 2020) as differentially expressed/methylated in at least one of the pairwise comparisons between islands. Further, 42 out of the 122 (34.4%) Denisovan-driven methylQTL targets were differentially methylated, and 7 (22.6%) and 149 (25.6%) of the Neanderthal-driven eQTL and methylQTL targets were differentially expressed/methylated. These observations highlight how both modern ancestry and archaic introgression can contribute to between-population differences in molecular phenotypes in the region.

**Figure 4.**
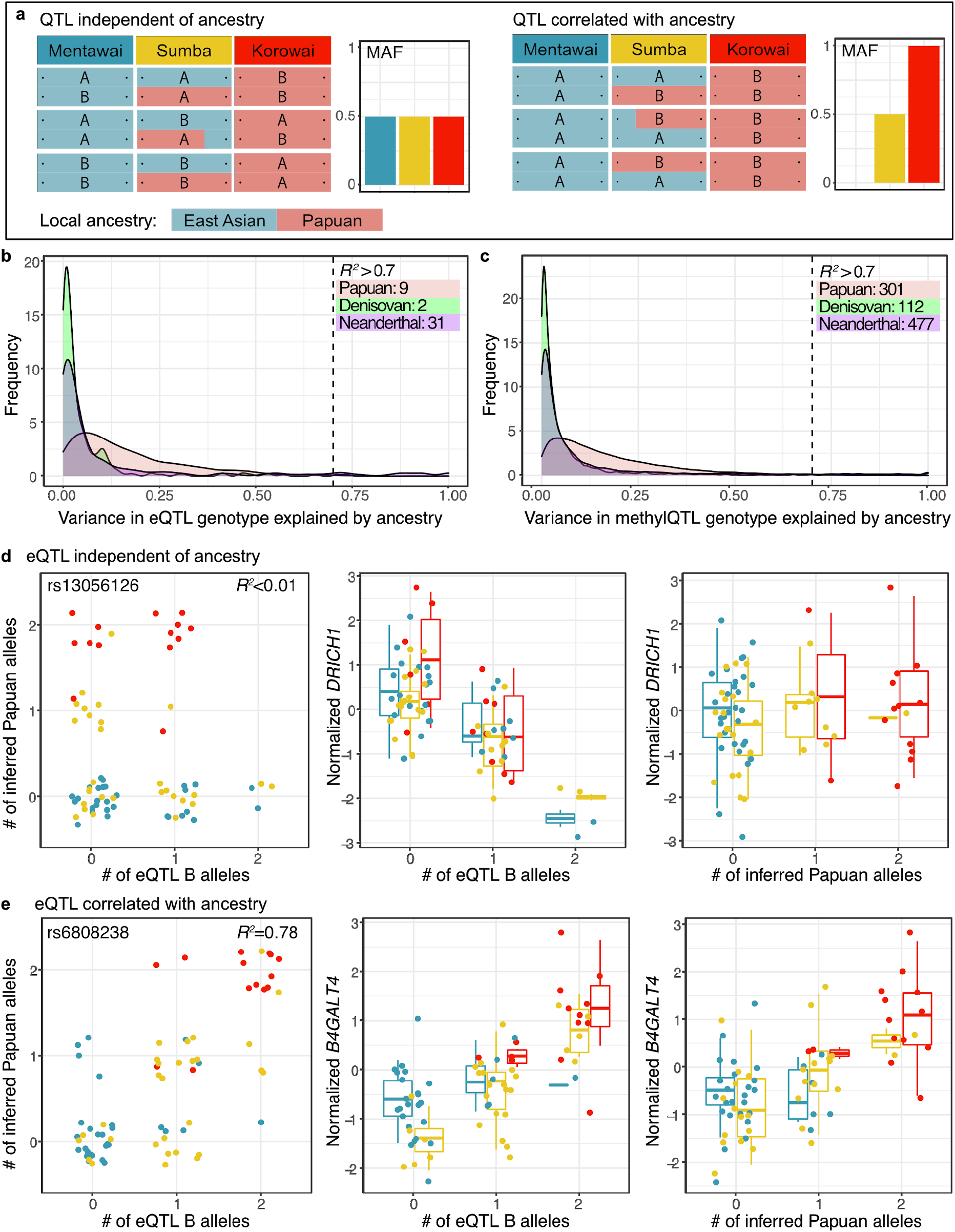
Integrating local ancestry inference at regulatory loci to detect QTLs driven by ancestry and archaic introgression. **a:** Schematic illustrations of variation in QTL genotype (A = major allele, B = minor allele) and local ancestry are shown across the two haplotypes in three individuals in three populations. In the first example, QTL genotype variation is independent of local ancestry, and allele frequencies are equal between populations. Thus, there is an expected correlation between the QTL genotype and the molecular trait, but not between ancestry and the trait. In the second example, QTL B allele closely segregates with the ancestry informative marker, and allele frequencies differ between populations. There is an expected correlation between the genotype and the molecular trait, as well as inferred ancestry and the trait. **b, c:** Linear regression between the numbers of QTL B alleles and numbers of inferred Papuan, Denisovan, and Neanderthal alleles reveal subsets of (**b**) eQTLs and (**c**) methylQTLs largely driven by modern LA and archaic introgression. The numbers of QTLs exceeding the *R^2^* threshold of 70% are indicated. **d:** An example of an eQTL independent of modern LA. **e:** An example of an eQTL highly correlated with modern LA. In **d** and **e**, the leftmost plot shows the correlation between the number of inferred Papuan alleles and eQTL B alleles. rs ID and *R^2^* are indicated. The center plot shows the effect of the eQTL B allele dosage on the normalized expression level of the target gene. The rightmost plot shows the effect of the inferred Papuan allele dosage on the target gene.

**Figure 5.**
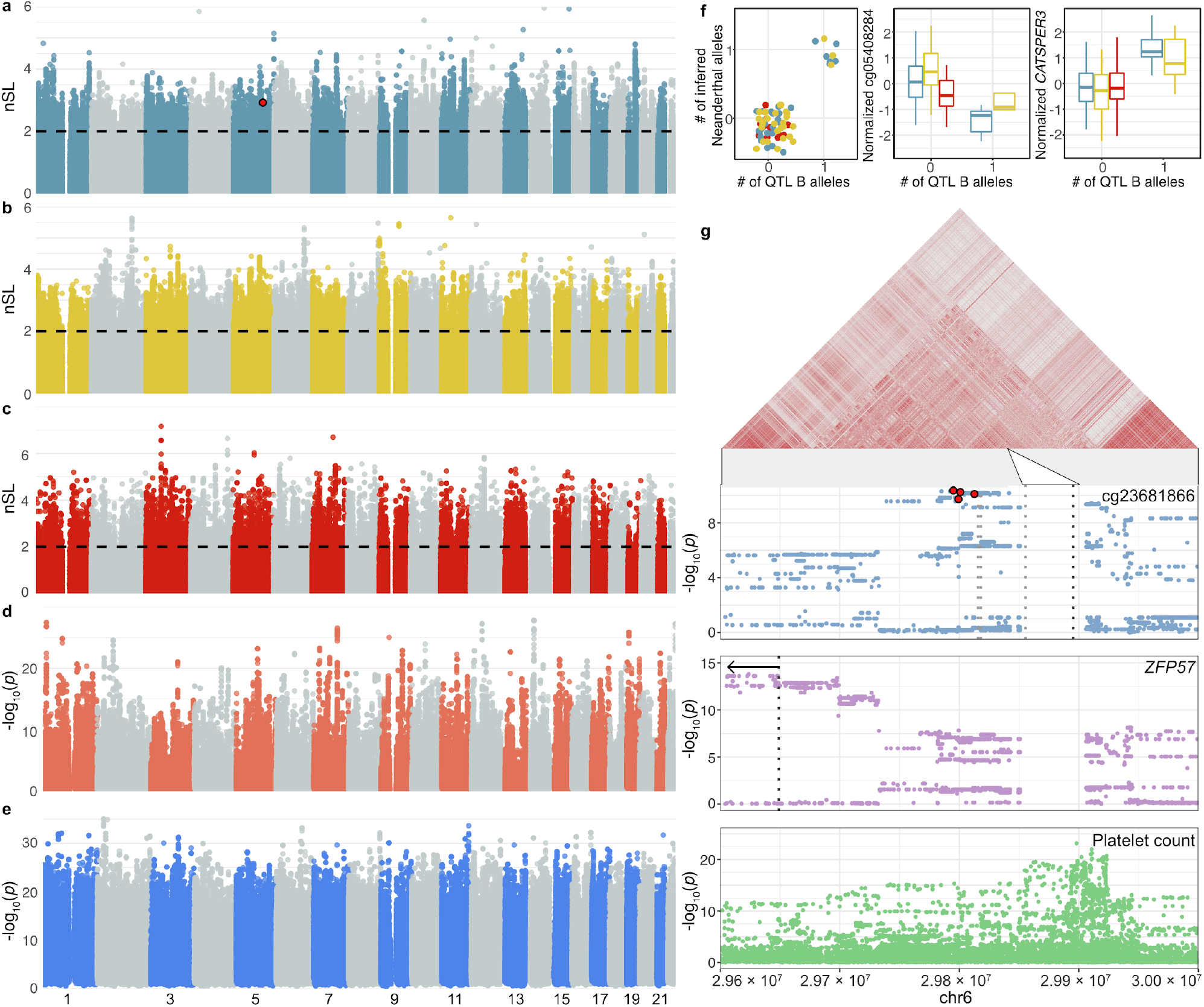
Subsets of LA and archaic introgression driven QTLs under positive selection or connected to complex traits. **a**, **b**, **c:** Manhattan plots of absolute nSL values across the autosomes in Mentawai (**a**), Sumba (**b**) and Korowai (**c**). nSL threshold of 2 is indicated. The QTL plotted in **f** is indicated in red in **a**. **d**, **e**: Manhattan plots of eQTLs (**d**) and methylQTLs (**e**). The x-axis shows positions along the autosomes and the y-axis shows the −log_10_(*p*-value) for each SNP. For each QTL SNP, −log_10_ of the smallest *p*-value is plotted. **f**: An example of a Neanderthal-driven QTL under positive selection Mentawai. The left-side plot shows the relationship between the QTL genotype and the number of inferred Neanderthal alleles. The center-plot shows the relationship between the methylQTL genotype and normalized CpG methylation. The right-side plot shows the relationship between the genotype and normalized *CATSPER3* expression. **g**: An example of a Denisovan-driven QTL that colocalizes with platelet count GWAS signal. Manhattan plots of the cg23681866 methylQTL, *ZFP57* eQTL, and platelet count GWAS. Dotted lines indicate CpG positions and *ZFP57* TSS. Three additional methylQTLs on the region colocalize with platelet count, the top SNPs of all four methylQTLs are indicated in red. All four methylVariants are nominally associated with *ZFP57*. Locations of the three additional methylQTL CpGs are indicated with light gray dotted lines. Indonesian LD patterns along this part of the genome are shown above.

### A subset of archaic ancestry driven QTLs are under positive selection in the region

We asked whether positive selection on ancestry informative regulatory variants may have contributed to the between-population variation in molecular phenotypes in the region. We used a haplotypebased nSL selection scan (Methods) to identify genomic regions that show signs of past selective sweeps and found 4.7%, 4.6%, and 5.0% of the genome to be under positive selection in Mentawai, Sumba and Korowai, respectively. We overlapped the ancestry driven QTLs with genomic regions with strong evidence of positive selection. While we detect no clear overrepresentation of ancestry-driven QTLs among these regions (Supplementary Table 4), we find individual QTLs that overlap them (Supplementary Table 7, Extended Data Figures 4, 5), including one Papuan-driven eQTL under selection in Mentawai, one Neanderthal-driven eQTL in Korowai and one in Mentawai, as well as Papuan-driven methylQTLs under selection in Mentawai (2), Sumba (2), and Korowai (12).

Moreover, we detect one Denisovan-driven methylQTL under selection in Korowai, associated with a CpG located on the promoter of *ZNF426*. Genetic variation associated with *ZNF426* and other KRAB-ZNF genes has previously been identified on candidate regions for positive selection in multiple human populations (Perdomo-Sabogal and Nowick 2019; Ávila-Arcos et al. 2020). Further, we identified 13, 6, and 3 Neanderthal-driven methylQTLs under selection in Mentawai, Sumba, and Korowai (Supplementary Table 5). For example, a Neanderthal-driven methylQTL under selection in Mentawai was also nominally associated (*p* = 2.596×10^−7^) with *CATSPER3* (Cation Channel Sperm Associated 3) expression, which was differentially expressed between Mentawai and Korowai, as well as Sumba and Korowai (Natri et al. 2020). Neanderthal variation in sodium channel genes was recently linked to increased pain sensitivity in modern humans (Zeberg et al. 2020). These findings indicate that positive selection on introgressed archaic variation may have shaped the patterns of these molecular traits, as well as their downstream functions, in Indonesia.

### Connecting regulatory variants to complex traits

GWAS colocalization analyses offer a way to connect regulatory variants to biological functions and complex traits. We tested for colocalization between the significant QTLs and 36 hematological traits using genome-wide summary statistics from a GWAS study on 173,480 European ancestry participants (Astle et al. 2016). We detected 30 and 614 unique eGenes and methylCpGs that colocalize with 34 and 36 traits, respectively (Supplementary Table 8). Notably, we find a Papuan-driven methylQTL that colocalizes with hemoglobin measurements, as well as four Denisovan-driven methylQTLs that colocalize with platelet count. We further examined these four methylQTLs to gain insight into possible mechanisms underlying the connection with platelet count (Supplementary Note 4, Supplementary Table 9). All four target CpGs are located near the HLA superlocus. While these methylQTLs do not significantly colocalize with eQTLs in our data, they are nominally associated with the nearby *ZFP57. ZFP57* is a transcriptional regulator known to have an important role in DNA methylation, epigenetic regulation and imprinting during development (X. Li et al. 2008). Expression of *ZFP57* is dependent on underlying genetic variation, and while the biology of *ZFP57* in adults is not well studied, its location within the HLA region points to a role in immunity, and it has been implicated as the causal gene connecting some GWAS variants to cancer and HIV/AIDS progression (Plant et al. 2014). Further multi-ethnic GWAS and functional studies are needed for the fine-mapping of causal variants underlying transcription.

## Discussion

Indonesia is the world’s fourth most populous country. Its people represent a key region that has been vastly understudied, one that is undergoing a rapid demographic and lifestyle shift giving rise to an expanding middle class, and where non-infectious, complex diseases are already contributing substantially to mortality and morbidity. This transition accelerates the need to understand the molecular underpinnings of complex disease, and in this context, our study adds to a growing literature demonstrating the importance of characterizing functional genomics within traditionally understudied populations (Mogil et al. 2018; Tehranchi et al. 2019).

We have explored the degree to which functional variation differs between Europeans and Indonesians, and more broadly, the problem of translating eQTL knowledge across populations. When comparing the eQTLs identified in this study to those identified previously in a European cohort (Lepik et al. 2017), only 45% have evidence of a shared causal variant. Additionally, we identified 166 genes that are eQTLs in Indonesians but have no evidence of an eQTL in Europeans, representing unique population-specific eQTLs. These eQTLs impact the expression of genes involved in clinically relevant biological processes, such as immunity and cancer progression.

Leveraging the unique cline of Asian and Papuan ancestry in Indonesia, we were also able to identify both eQTLs and methylQTLs driven by local ancestry or introgression from archaic hominin species. In an attempt to identify the forces shaping QTLs driven by LA or archaic alleles, we integrated the QTL results with signals of recent positive selection in the three Indonesian study populations, possibly indicative of local adaptation. While limited in number, we do indeed identify such QTLs that are also under positive selection. At least anecdotally, this demonstrates how local selective pressures can act upon unique non-coding variation altering molecular traits and presumably endpoint phenotypes as well. This is further bolstered by our identification of QTLs co-localizing with GWAS traits. Taken together these data demonstrate how population-level genetic structure can drive differences in functional variation that contributes to complex traits.

We believe the modest overlap of GWAS hits with the population-specific QTLs represents a nontrivial challenge in the field of functional genomics: how do we connect population-specific functional variation to loci associated with complex traits identified in European populations? From a practical perspective, we do not anticipate a robust expansion of traditional GWAS studies being carried out in understudied populations. To this end, the field will need to move away from simple intersections of GWAS and QTL hits, which rely upon shared LD structure, and instead integrate genetic variation, GWAS results, context-specific multi-omics (in simulated or actual disease states and in a range of cell types), and robust functional validations to define common sets of regulatory elements that contribute to disease and are shared across populations.

## Materials and methods

### Ethical approvals

All samples in this study have been previously reported (Natri et al. 2020). Samples were collected by HS and an Indonesian team from the Eijkman Institute for Molecular Biology, Jakarta, Indonesia, with the assistance of Indonesian Public Health clinic staff. All collections followed protocols for the protection of human subjects established by institutional review boards at the Eijkman Institute (EIREC #90 and EIREC #126) and the University of Melbourne (Human Ethics Sub-Committee approval 1851639.1). All individuals gave written informed consent for participation in the study. Permission to conduct research in Indonesia was granted by the Indonesian Institute of Sciences and by the Ministry for Research, Technology and Higher Education.

### Data acquisition

Here we report two new genomic datasets: 1) 42 samples genotyped using the Illumina Infinium Omni2.5-8 v1.3 BeadChip array, including 5 Korowai samples from New Guinea, 18 samples from Mentawai, western Indonesia, and 19 samples from Sumba, eastern Indonesia; 2) complete genomes for 70 samples sequenced to an expected mean depth of 30x, including 11 Korowai, 30 Mentawai, and 29 Sumba samples.

### Whole genome data processing

The newly generated genome sequences were processed closely following the protocol described in (Jacobs et al. 2019) with the resources of the University of Tartu High Performance Computing Center (University of Tartu 2020). Briefly, we first aligned the reads to the ‘decoy’ version of the GRCh37 human reference sequence (hs37d5). After alignment, and keeping only properly paired reads that mapped to the same chromosome, the autosomal sequencing depth across the samples used in downstream analyses was as follows: min = 31.5x, Q1 = 35.3x, median = 36x, Q3 = 36.5x, max = 39.5x. Base-calling was undertaken using GATK best practices (Poplin et al., n.d.; Auwera et al. 2013). Following the generation of per-sample gVCF files with GATK4 HaplotypeCaller, single sample gVCFs were combined into multisample files using CombineGVCFs, and joint genotyping was performed using GATK4 GenotypeGVCFs, outputting all sites to a multisample VCF. To maximize the SNP discovery and phasing power, approximately 900 complete genomes were used in a multisample calling pipeline. In addition to the newly generated genomes, these included complete genome sequences from SGDP (Mallick et al. 2016) and IGDP (Jacobs et al. 2019) projects, Malaspinas et al. (Malaspinas et al. 2016), Vernot et al. (Vernot et al. 2016), Lan et al. (Lan et al. 2017), and the HiSeqX Diversity Cohort of Polaris project (https://github.com/Illumina/Polaris), as well as approximately 100 unpublished genome sequences from Estonia and Papua. SNP calling was performed on the combined dataset, with published genomes analyzed from raw reads exactly as for the new sequence data. Using bcftools v1.9 (H. Li 2011), the following filters were applied to each genotype call in multisample VCF files: base depth (DP) ≥8x and ≤400x, and genotype quality (GQ) ≥30. Only biallelic SNPs and invariable reference sites were kept.

The published data included 7 Korowai and 10 Mentawai samples, however, two first-degree relatives (MTW024 and MTW066) were excluded from further analysis (Jacobs et al. 2019). Our final WGS dataset therefore included 84 samples from three target groups: 17 Korowai, 38 Mentawai, and 29 Sumba.

Next, modern human multisample VCF files were merged with two archaic individuals: Denisovan (Meyer et al. 2012) and Neanderthal (Prüfer et al. 2014). Positions with missing or low-quality calls (marked as ‘LowQual’ in the original archaic VCF files) in one of the archaic samples were excluded during the merging procedure. We kept only sites that had high-quality variant calls in at least 99% of samples in the combined modern/archaic dataset. Applying this 99% call-rate filter yielded a total of 52,443,217 SNPs. However, we removed sites within segmental duplications, repeats, and low complexity regions, thus retaining 49,374,343 SNPs. These masks were downloaded from the UCSC and Broad Institute genome resources: http://hgdownload.soe.ucsc.edu/goldenPath/hg19/database/genomicSuperDups.txt.gz http://software.broadinstitute.org/software/genomestrip/node_ReferenceMetadata.html Phasing was performed with Eagle v2.4 (Loh, Palamara, and Price 2016). Because our final dataset included complete genomes from very diverse human populations together with a large number of local Island Southeast Asian and Papuan groups, we did not use any reference datasets to avoid potential phasing bias.

### Genotype array data processing

Array data for 42 Indonesian and Papuan individuals was processed in PLINK v1.9 (Chang et al. 2015). The average missing rate per person in the raw dataset was around 0.45% (min 0.27%, max 2.5%); 2,194,297 autosomal positions were kept after excluding SNPs with more than 5% of missing data.

Array data was imputed with Beagle v5.1 (Browning, Zhou, and Browning 2018) using complete genome sequences as a reference. Two imputation reference panels were generated. For the imputation of 18 Mentawai samples, we applied a reference panel that included 97 complete genome sequences from western Indonesia (Bali, Borneo, Java, Mentawai, Nias, Sulawesi, Sumatra), Philippines and Taiwan. For the imputation of 24 Korowai and Sumba samples, we applied a reference panel made of 249 complete genomes sequence from eastern Indonesia (Alor, Flores, Kei, Lembata, Sumba, Tanimbar) and Papua (Bougainville, New Britain, New Guinea, including Korowai, and New Ireland).

Variant sites were filtered using bcftools and VCFtools (Danecek et al. 2011) to retain only high quality imputed sites with dosage *R^2^* >0.95 (estimated squared correlation between the estimated allele dose and the true allele dose, *DR^2^*). These positions were extracted from the complete genomes from Korowai, Mentawai and Sumba (N = 84) to produce a new combined SNP set made of imputed and WGS data. These data were filtered to retain SNPs with a proportion of missing data < 0.3 and minor allele frequency (MAF) >0.05, which resulted in 4,077,164 variants. Imputed genotypes were further filtered to retain genotypes with genotype probability (GP) >0.90.

### Local ancestry inference

We used ChromoPainter v2 CP, (Lawson et al. 2012) to perform local ancestry (LA) inference and detect Asian and Papuan ancestry in all published and newly generated complete genomes from Korowai (N = 17), Mentawai (N = 38) and Sumba (N = 29). This method relies on phased haplotype data and describes each individual recipient chromosome as a mixture of genetic blocks from the set of predefined donor individuals.

First, East Asian and Papuan reference panels were generated to assign local genomic ancestry in target samples. We selected unadmixed East Asian and Papuan samples by running ADMIXTURE v1.3 (Alexander et al. 2009) at K = 3 using all available East and Southeast Asian, European and Papuan samples from the combined WGS dataset. For the East Asian reference panel, we kept only Asian samples (N = 102) with less than 0.05% of non-East Asian ancestry. For the Papuan reference panel, we kept only Papuan samples (N = 63) with less than 0.05% of non-Papuan ancestry and excluded all Korowai samples. To balance the sample size of the two reference panels, we randomly selected 63 East Asian samples from the unadmixed reference dataset.

Next, we painted each of 84 target genomes individually using the East Asian and Papuan reference panels as donors. We used the following protocol:

1. The initial CP run was performed with 10 EM steps to estimate prior copying probabilities for each individual and chromosome separately.
2. Estimated prior copying probabilities were averaged across the genome for each individual. The main CP run was performed with a recombination scaling constant and global mutation probability from the first step, and genome-wide average prior copying probability.
3. Either East Asian or Papuan ancestry was then assigned to individual SNPs using a probability threshold of 0.85. Unknown ancestry was assigned to SNPs with intermediate copying probability.

### Identifying archaic Denisovan introgression

We defined the high confidence Denisovan archaic haplotypes as outlined previously (Jacobs et al. 2019), but using a larger group of sub-Saharan African individuals (61 sub-Saharan Africans in total, Supplementary Table 3) For each individual, we started with Denisovan introgressed haplotypes as inferred by CP; then filtered out those that did not overlap (by >0.001%) the Denisovan introgressed haplotypes as inferred by a previously published hidden Markov model (HMM) (Jacobs et al. 2019); then filtered out those that did not overlap (by >0.001%) archaic introgressed haplotypes inferred by the another HMM approach (Skov et al. 2018); and finally filtered out any of the remaining haplotypes that did overlap (by >0.001%) Neanderthal introgressed haplotypes as inferred by CP. We then annotated each SNP found in several target sample groups (i.e., monomorphic SNPs in that group are skipped, as are any that are masked out by the alignability/gap mask) according to how often the REF/ALT state appears on an inferred high confidence Denisovan introgressed haplotype in that group. This was done for three separate groups: 1) all Korowai individuals, 2) all Korowai individuals, Sumba individuals and those Mentawai individuals who are from the new dataset, and 3) all individuals in the ‘Papuan’ continental group, which includes all Papuans and Melanesians except Baining. An analogous process was used to annotate Neanderthal ancestry SNPs, beginning instead with Neanderthal introgressed haplotypes inferred by CP before requiring intersection with Neanderthal introgressed haplotypes inferred by the HMM and archaic haplotypes inferred by HMM_Archaic_ and removing those intersecting CP Denisovan haplotypes.

### DNA methylation data processing

DNA methylation data were processed as previously described (Natri et al. 2020) using *minfi* v1.30.0 (Aryee et al. 2014) The two arrays were combined and preprocessed to correct for array background signal. Signal strength across all probes was evaluated and probes with signal *p* < 0.01 in >75% of samples were retained. To avoid potential spurious signals due to differences in probe hybridization affinity, we discarded 6,072 probes overlapping known SNPs segregating in any of the study populations based on previously published genotype data (Jacobs et al. 2019). The final number of probes retained was 859,404. Subset-quantile Within Array Normalization (SWAN) was carried out using the ‘preprocessSWAN’ function (Maksimovic, Gordon, and Oshlack 2012). Methylated and unmethylated signals were quantile normalized using *lumi* v2.36.0 (Du, Kibbe, and Lin 2008).

### Gene expression data processing

RNA sequence data were processed as in (Natri et al. 2020) FASTQ read files underwent quality control with FastQC v0.11.5 (Andrews 2010), and leading and trailing bases below a Phred score of 20 were removed using Trimmomatic v0.36 (Bolger, Lohse, and Usadel 2014). Reads were aligned to the human genome (GRCh38 Ensembl release 90: August 2017) with STAR v2.5.3a (Dobin et al. 2013) and a two-pass alignment mode. Read counts were quantified with featureCounts v1.5.3 (Liao, Smyth, and Shi 2014) against a subset of GENCODE basic (release 27) annotations that included only transcripts with support levels 1–3. Coordinates were converted to hg19 with the R package *liftOver* v1.8.0 (Bioconductor Package Maintainer 2020). Gene expression data were filtered to retain 12,539 genes with FPKM >0.1 and read count of >6 in at least 50 samples. The distributions of FPKM in each sample and gene were transformed into the quantiles of the standard normal distribution.

### Accounting for population structure and non-genetic sources of variation in the QTL analyses

Principal Component Analysis (PCA) of the genotype data was carried out using the R package *SNPRelate* v1.18.1 (Zheng et al. 2012). Five genotype PCs were included as covariates in QTL analyses to account for population structure. We used a probabilistic estimation of expression residuals (PEER, (Stegle et al. 2012)) to infer hidden sources of variation in expression and methylation data. These latent factors were used as surrogate variables for unknown technical batch effects and included as covariates the QTL analyses. 29 hidden factors (25% of the number of samples) were included in models, as recommended in (Stegle et al. 2012).

### eQTL and methylQTL analyses

Variant effects on gene expression and CpG methylation were identified by linear regression as implemented in QTLtools (Delaneau et al. 2016). Genotype, gene expression and methylation data were available for 115 individuals: 48 Mentawai, 48 Sumba, and 19 Korowai (Supplementary Tables 1 and 2). Variants within 1Mb of the gene/CpG under investigation were considered for testing. *p*-values of top-associations adjusted for the number of variants tested in *cis* were obtained using 10,000 permutations. False discovery rate (FDR) adjusted *p*-values were calculated to adjust for multiple phenotypes tested. Significant associations were selected using an FDR adjusted *p*-value threshold of 0.01. Nominal *p*-values for all sites within the *cis*-window were obtained using the QTLtools nominal pass. QTL power calculations were carried out using the R package *powerEQTL* v0.1.7 (Dong et al. 2017).

### Variant annotation and variant set enrichment analyses

To understand the genomic context of the putative eQTLs and methylQTLs, top-SNPs from the permutation-based analyses and the target CpGs of methylQTLs were annotated using the R package *annotatr* v1.10.0 (Cavalcante and Sartor 2017). Genic annotations (1-5Kb upstream of the TSS, the promoter (< 1Kb upstream of the TSS), 5’ UTR, first exons, exons, introns, coding sequences (CDS), 3’ UTR, and intergenic regions) were obtained using the *TxDb.Hsapiens.UCSC.hg19.knownGene* R package v3.2.2 (Carlson & Bioconductor Package Maintainer 2015), CpG annotations using the *AnnotationHub* R package v2.16.1 (Morgan & Shepherd 2020), and enhancer annotations from FANTOM5 (Andersson et al. 2014)

We tested for the enrichment of the eQTL and methylQTL variants among genomic features using the *VSE* R package v0.99 (Ahmed et al. 2017). A null-distribution was constructed based on 100 matched random variant sets. Consolidated ChIP-seq peaks for histone marks derived from primary mononuclear cells from peripheral blood were downloaded from the NIH Epigenomics Roadmap FTP site (Chadwick 2012). Additionally, annotations for DNaseI hypersensitivity peaks and histone marks for K562 and GM12878 cell lines were downloaded from the ENCODE portal (Davis et al. 2018).

We tested for the overrepresentation of the Indonesian population-specific eGenes among GO terms and canonical pathways using *clusterProfiler* 3.14.3 (Yu et al. 2012).

### eQTL-methylQTL colocalization analysis

We used a Bayesian test, as implemented in the R package *coloc* v4 (Giambartolomei et al. 2014; Wallace 2020), to assess the probability of co-localization of methylQTL and eQTL signals between 3,057 pairs of CpGs and genes. We used masking to allow for multiple causal loci for each trait. Masking implemented in *coloc* allows for multiple causal variants per trait with the assumption that if multiple causal variants exist for any individual trait, they are in linkage equilibrium. All SNPs independently associated within a dataset were identified with finemap.signals(). For the pairs of CpGs and genes with multiple signals, colocalization analysis was performed for each pair of signals, restricting the search space to SNPs not in LD with any-but-one of each signal SNP. The *p*-value threshold for calling a signal was set to 1×10^−6^, and the maximum *r^2^* between two SNPs for them to be considered independent was 0.01.

Pairs with the posterior probability for a CCV > 0.8 and the ratio of the posterior probability for a CCV and different causal variants (DCV) CCV/DCV >5 were considered to show strong evidence of colocalization. As the posterior probability for colocalization is dependent on the prior probability, we used the *coloc* post-hoc sensitivity analysis to determine the range of prior probabilities (1.0×10^−8^ to 1.0×10^−4^) for which colocalization is supported. Pairs passing the colocalization threshold with a range of ppCCV values from <1.0×10^−6^ to 1.0×10^−4^ (lower bound of ppCCV below 1.0×10^−6^) were considered as showing robust support for colocalization.

### Colocalization with European eQTLs and blood trait GWAS loci

Similarly to eQTL-methylQTL colocalization, we used *coloc* v4 to test for colocalization between 3,300 permutation-based eQTLs detected here with an FDR-adjusted *p* < 0.10 and data from the (Lepik et al. 2017) blood dataset (N = 491). Lepik et al. 2017 eQTL summary statistics were obtained using the EBI eQTL catalog API (Kerimov et al., n.d.). The methods used to call the eQTLs in the EBI eQTL catalog are comparable to the methods used in this study. Out of the 3,300 genes selected for testing, 3,049 were present in the European data and had shared variant with the Indonesian data. We identified colocalized genes with the threshold CCV > 0.8 and a ratio CCV/DCV > 5. To identify genes that do not show support for colocalization even with a relaxed threshold, we used a threshold of CCV > 0.5 and CCV/DCV > 2. To connect the QTLs detected here to blood traits, we tested for colocalization between the FDR-significant permutation-based QTLs and 36 hematological traits using genome-wide summary statistics from (Astle et al. 2016). GWAS summary statistics were downloaded from the GWAS catalog (MacArthur et al. 2017). As no LD information was available, these colocalization analyses were carried out without allowing for multiple causal variants.

### Selection scanning

We performed an nSL scan (Ferrer-Admetlla et al. 2014), as implemented in Selscan v1.2.0 (Szpiech and Hernandez 2014). This test identifies ongoing positive selection in the genome by looking for the tracts of extended haplotype homozygosity and is capable of identifying both sweeps from standing variation and incomplete sweeps. To identify the traces of positive selection in three target populations, we used our combined dataset of WGS and imputed genotyping array data represented by approximately 4M SNPs. The following Selscan parameters were used: maximum allowed gap between loci of 50 Kb, the gap scale parameter of 5 Kb, maximum extent of haplotype homozygosity decay curve of 1,333 loci (approximately 1Mb given the obtained SNP density). Raw nSL results were normalized with Selscan’s *norm* package in 50Kb non-overlapping genomic windows using ten allele frequency bins. Windows with less than 21 SNPs were discarded. The proportion of absolute nSL scores > 2 in each 50Kb genomic window was used as a test statistic. Windows with a proportion of SNPs with an absolute nSL > 2 of 30% were considered to be outliers and showing evidence of past positive selection.

### Identifying eQTL effects driven by local ancestry

We calculated the variance explained by modern LA in the genotype of each significant (FDR-*p* < 0.01) permutation-based eQTL and methylQTL as in (Gay et al. 2019). For each eVariant and methylVariant, we fit the linear model *V* = *α* * *PAP* + *β*, where V is the genotype vector (number of QTL B alleles), and PAP is the LA covariate, representing the number of alleles assigned to the Papuan population. This analysis was carried out using the 73 WGS (30 Mentawai, 29 Sumba, 14 Korowai) samples included in the LA inference. Variants with an absolute *R^2^* > 0.7 were considered to exhibit a high correlation with LA. Similarly, we calculated the variance explained by archaic Denisovan and Neanderthal ancestry.

### Data availability

All genotype data, RNA sequencing reads, and Illumina Epic iDat files are available through the Data Access Committee of the official data repository at the European Genome-phenome Archive (EGA; https://www.ebi.ac.uk/ega/home). Illumina Omni 2.5M genotyping array data are deposited in study EGAS00001003670 and the whole genome data in study EGAS00001003654. The RNA sequencing data are deposited in study EGAS00001003671 and the methylation data are deposited in study EGAS00001003653. Matrices of unfiltered read counts (doi:10.26188/5d12023f77da8) and M-values (doi:10.26188/5d13fb401e305) for all samples, including replicates, are freely available on Figshare (https://figshare.com). Supplementary Files 3 and 4 containing the nominal eQTL statistics for all tested SNP-gene pairs and the nominal methylQTL statistics for the all SNPs tested for the targets of the permutation-significant methylQTLs are available on Figshare with doi:10.26188/12871007 and doi:10.26188/12869969, respectively.

## Acknowledgments

We especially thank all of our study participants. This study was supported by a Royal Society of New Zealand Marsden Grant 17-MAU-040 to MPC and IGR and by the ASU Center for Evolution and Medicine and the Marcia and Frank Carlucci Charitable Foundation postdoctoral award from the Prevent Cancer Foundation to HMN. IGR, GH and MM were partially supported by EU Horizon 2020 Grant 810645; MM and LS by European Regional Development Fund Projects 2014-2020.4.01.16-0030 and 2014-2020.4.01.15-0012 and by the Estonian Research Council grant PUT (PRG243).

## Extended Data

**Extended Data Figure 1.**
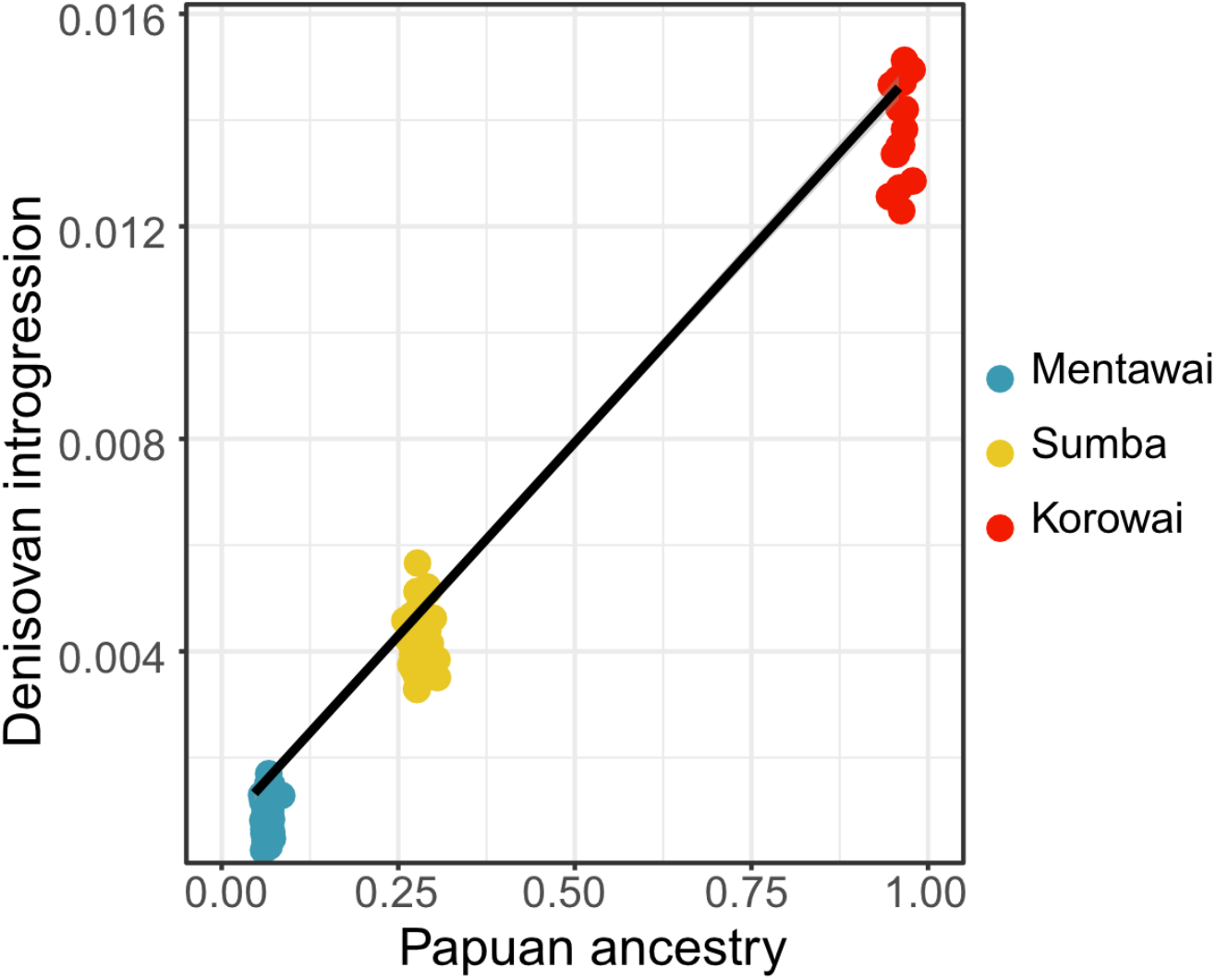
Proportions of inferred Papuan ancestry and Denisovan introgression are highly correlated (Pearson’s correlation coefficient 0.995).

**Extended Data Figure 2.**
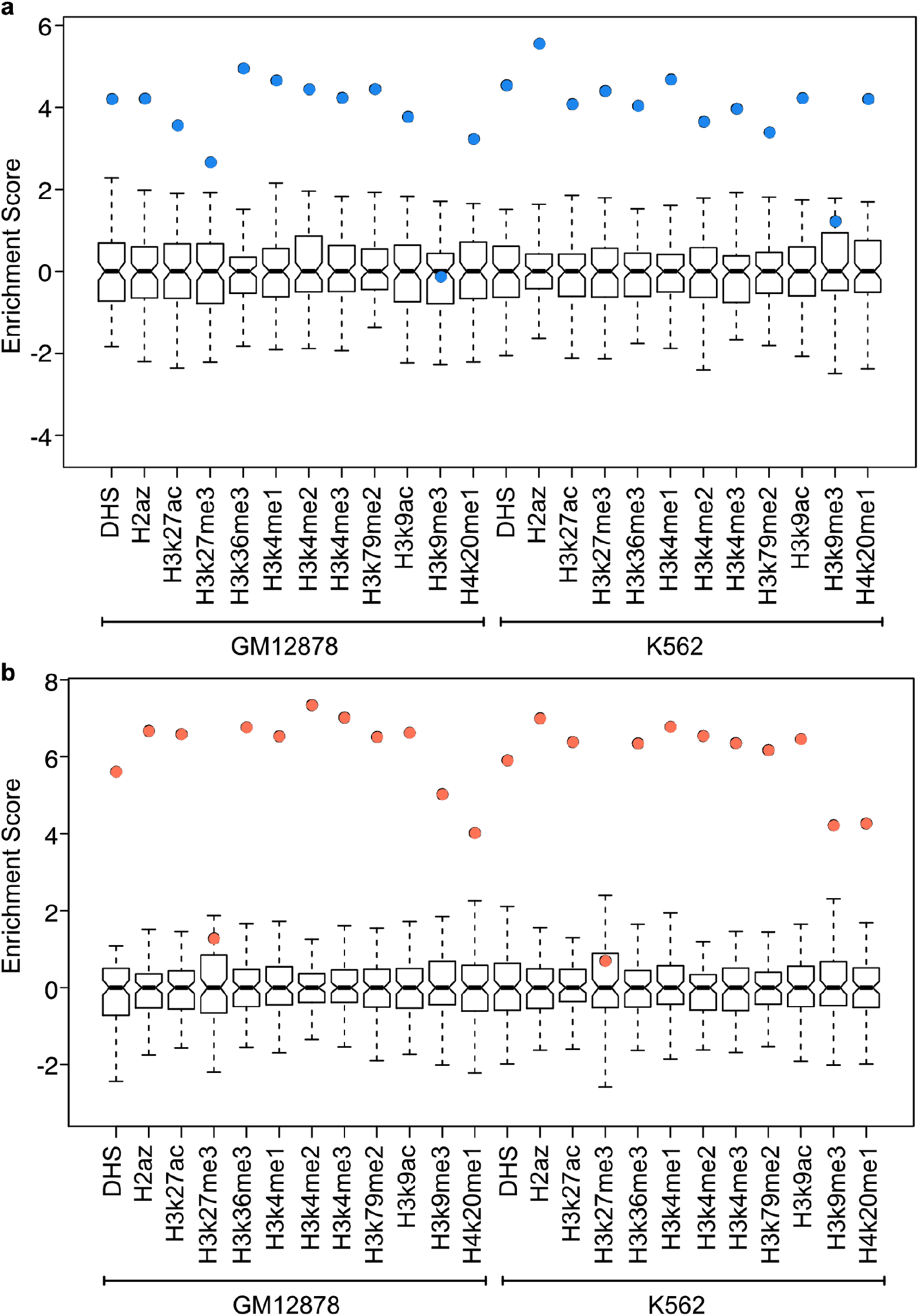
Enrichment of methylQTLs (**a**) and eQTLs (**b**) among DNase hypersensitive sites (DHS) and histone marks in ENCODE GM12878 and K562 cell lines.

**Extended Data Figure 3.**
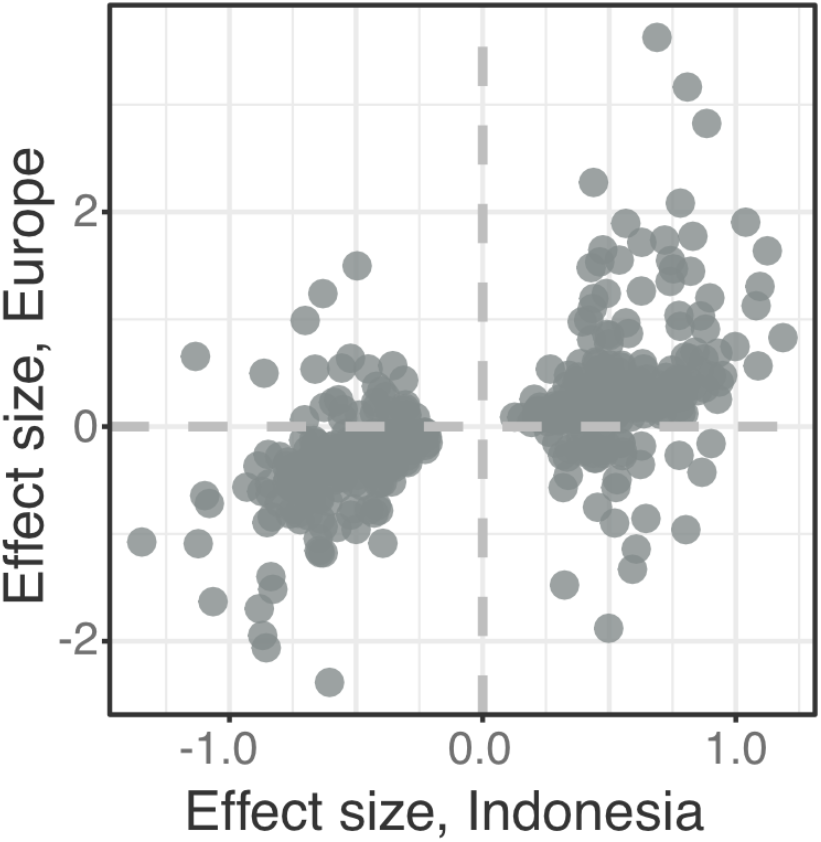
Effect sizes of colocalized eQTLs in the Indonesian and European datasets.

**Extended Data Figure 4.**
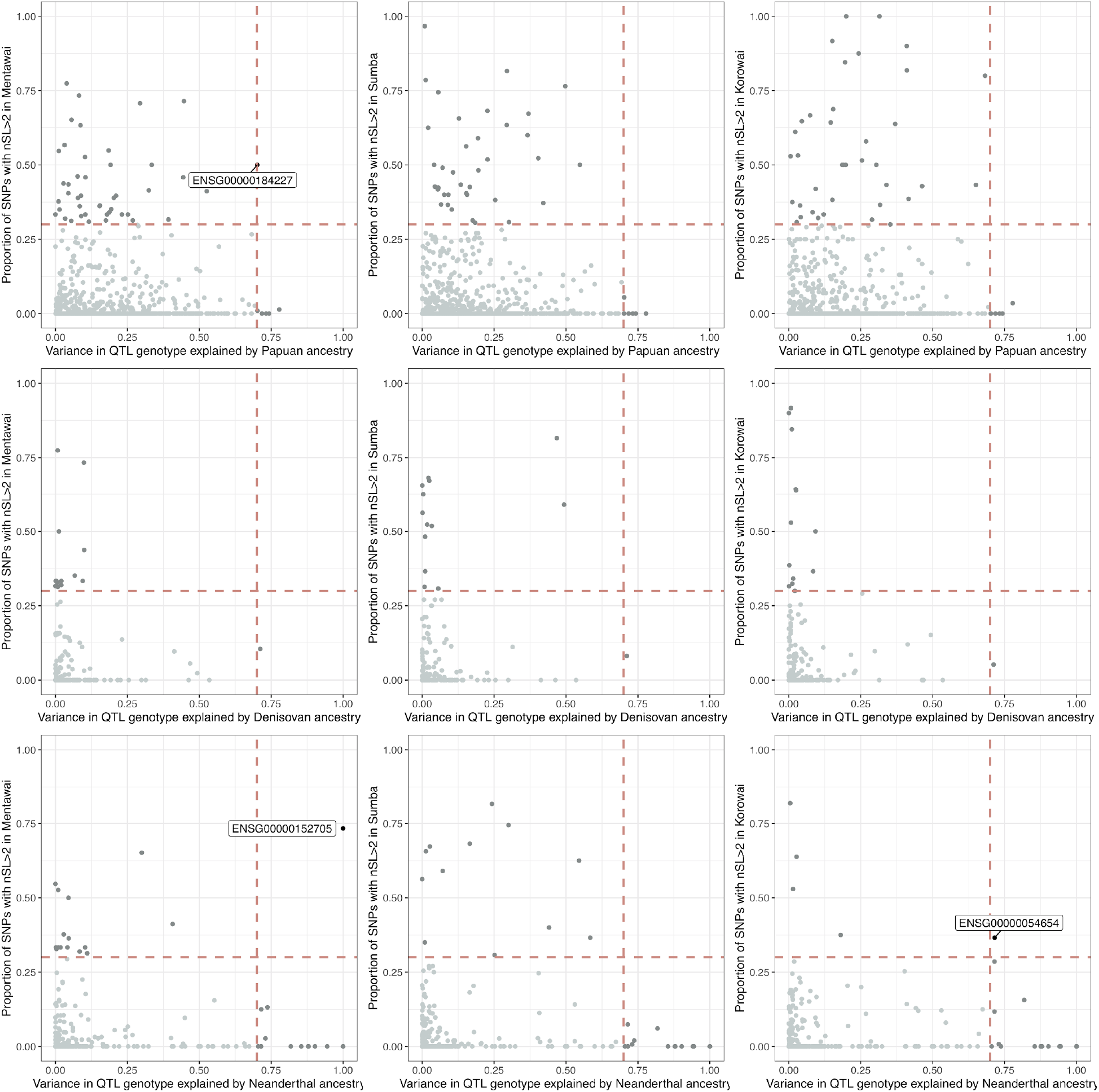
Modern ancestry and archaic introgression driven eQTLs overlapping genomic windows that show evidence of recent positive selection in each of the three study populations. Variance in QTL genotype explained (*R^2^*) is shown on the x-axis of each plot. Variants with *R^2^* > 0.7 were considered to be highly correlated with ancestry (vertical line). Proportion of positions within 50Kb windows that show an nSL > 2 is shown on the y-axis. Genomic windows with this proportion >0.3 were considered to be showing evidence of positive selection (horizontal line). The target genes of eQTLs showing both a significant correlation with ancestry and evidence of selection are labeled.

**Extended Data Figure 5.**
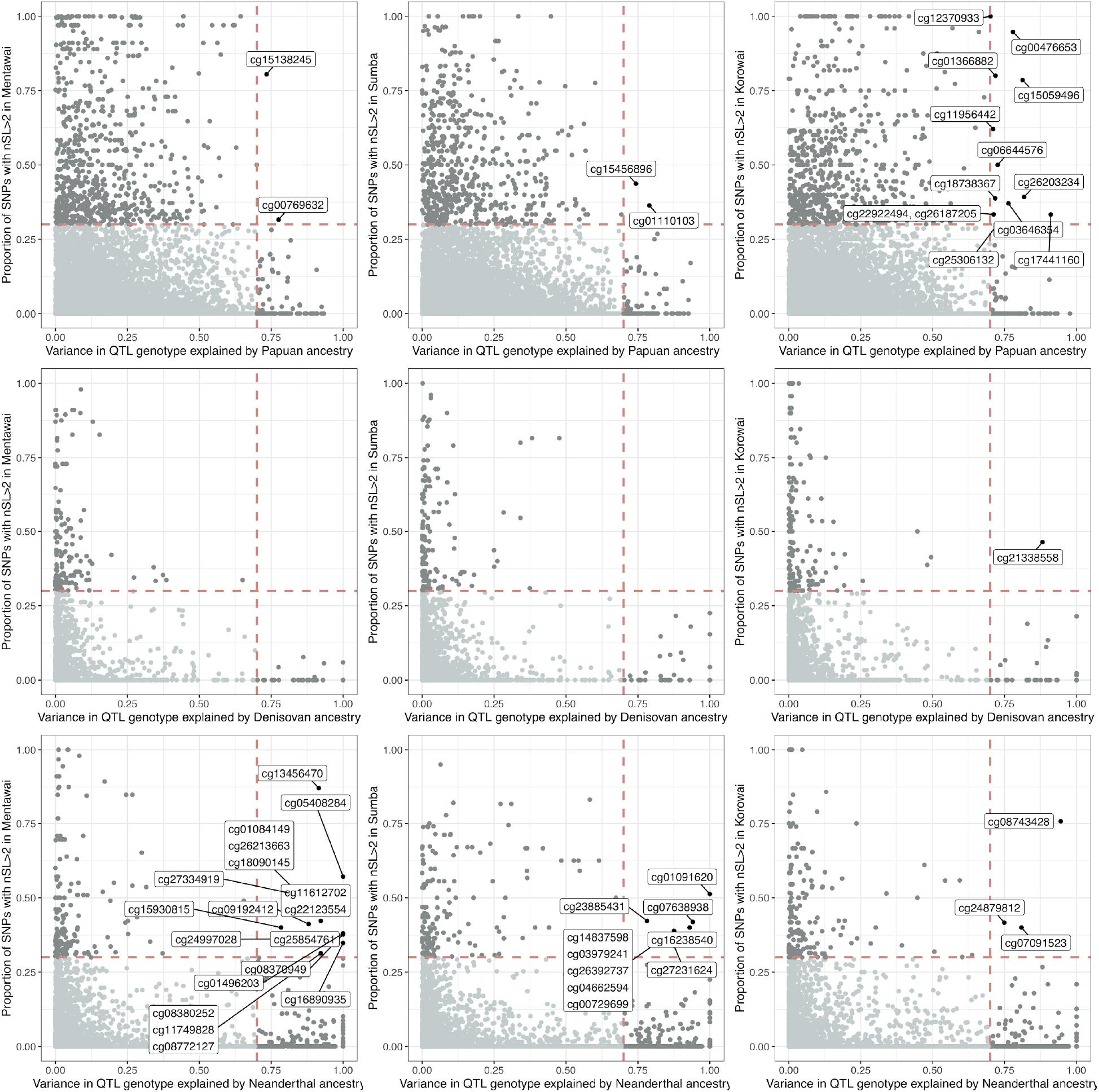
Modern ancestry and archaic introgression driven methylQTLs overlapping genomic windows that show evidence of recent positive selection in each of the three study populations. Variance in QTL genotype explained (*R^2^*) is shown on the x-axis of each plot. Variants with *R^2^* > 0.7 were considered to be highly correlated with ancestry (vertical line). Proportion of positions within 50Kb windows that show an nSL > 2 is shown on the y-axis. Genomic windows with this proportion >0.3 were considered to be showing evidence of positive selection (horizontal line). The target CpGs of methylQTLs showing both a significant correlation with ancestry and evidence of selection are labeled.

## Supplementary Information

### Supplementary_information.doc

- Supplementary Notes 1-4, Supplementary Figures 1-6

### Supplementary Tables 1-10

- Supplementary Table 1: Sample information.
- Supplementary Table 2: Sequencing batch information for RNAseq samples.
- Supplementary Table 3: Sub-Saharan African samples used in archaic introgression inference.
- Supplementary Table 4: Indonesian eGenes (FDR-*p* < 0.1) that do not colocalize with and are not eGenes (nominal p < 5×10^−4^) in the European data. Columns: Gene ID, gene name, European top SNP, European nominal *p*-value.
- Supplementary Table 5: GO and KEGG enrichment analysis on the 166 Indonesia-specific eGenes.
- Supplementary Table 6: Enrichment analysis of ancestry-driven QTLs among genomic regions under positive selection.
- Supplementary Table 7: Ancestry-driven QTLs overlapping genomic regions under positive selection.
- Supplementary Table 8: methylQTL-GWAS and eQTL-GWAS colocalization summary.
- Supplementary Table 9: Denisovan-driven methylQTLs that colocalize with platelet count GWAS.
- Supplementary Table 10: Megablast results for the four Denisovan/platelet count CpGs.

### Supplementary Files

- Supplementary File 1: Permutation-significant eQTLs. Columns: target, chromosome, target start, N of tested SNPs, top-SNP distance to the target, rsID, top-SNP position, slope, nominal *p*, FDR-*p*
- Supplementary File 2: Permutation-significant methylQTLs. Columns: target, chromosome, target position, N of tested SNPs, top-SNPdistance to the target, rsID, top-SNP position, slope, nominal *p*, FDR-*p*
- Supplementary File 3: Nominal eQTL statistics. Columns: target, target chromosome, target start, N of tested SNPs, SNP distance to the target, rsID, SNP position, nominal *p*, slope. The file is available on Figshare with DOI 10.26188/12871007.
- Supplementary File 4: Nominal statistics for permutation-significant methylQTLs. Columns: target, target chromosome, target start, N of tested SNPs, SNP distance to the target, rsID, SNP position, nominal *p*, slope. The file is available on Figshare with DOI 10.26188/12869969.
- Supplementary File 5: eQTL-methylQTL colocalization results for robust colocalized pairs. Columns: Target CpG, Target gene, number of tested SNPs, Tag SNP 1 and Tag SNP 2 (testing between all independent signals), PP0, PP1, PP2, PP3, PP4, PP4/PP3, lower bound of prior probability for colocalization (p12) that passes the threshold.
- Supplementary File 6: European colocalization results, significant genes. Columns: Target gene, number of tested SNPs, posterior probabilities for no association in either trait (PP0), association in trait 1 but not in trait 2 (PP1), association in trait 2 but not in trait 1 (PP2), association in both traits, different SNPs (PP3), and association in both traits, shared causal SNP (PP4), PP4/PP3, lower bound of prior probability for colocalization (p12) that passes the threshold.
- Supplementary Files 7: eQTL-LAI correlation results for Papuan ancestry. Columns: chr, pos, *R^2^*, *p* target
- Supplementary Files 8: eQTL-LAI correlation results for Denisovan introgression. Columns: chr, pos, *R^2^, p*, target
- Supplementary Files 9: eQTL-LAI correlation results for Neanderthal introgression Columns: chr, pos, *R^2^, p*, target
- Supplementary Files 10: methylQTL-LAI correlation results for Papuan ancestry. Columns: chr, pos, *R^2^*, *p*, target
- Supplementary Files 11: methylQTL-LAI correlation results for Denisovan introgression. Columns: chr, pos, *R^2^*, *p*, target
- Supplementary Files 12: methylQTL-LAI correlation results for Neanderthal introgression Columns: chr, pos, *R^2^*, *p*, target

