## Supplemental_materials for "Genetic architecture of gene regulation in Indonesian populations identifies QTLs associated with local ancestry and archaic introgression": Supplementary Information.docx

**Supplementary Note 1.** ***Colocalized cis-eQTLs and cis-methylQTLs indicate shared causal variants.*** We integrated the methylQTL and eQTL calls to gain insight into how genetic regulation of CpG methylation may contribute to the regulation of gene expression. 1,140 of the unique permutation significant eVariants were also nominally associated (nominal *p*<1×10^-7^) with the methylation of at least one CpG, and 2,015 of the unique permutation-significant methylVariants were also nominally associated with the expression of at least one gene, suggesting that a substantial number of causal eVariants may also be causal methylVariants, and vice versa. This overlap corresponds to 4,639 CpG-gene combinations potentially harboring a common causal variant (CCV). We tested for colocalization between these pairs of CpGs and genes using a Bayesian method as implemented in *coloc* v4. Among the tested pairs, we detected 720 (15.5%) eQTL-methylQTL pairs that showed robust support for colocalization with a wide range of prior probabilities for a common causal variant (Methods, Figure S1).

We explored the direction of the effects of top-SNPs associated with the 720 CpG and gene pairs that exhibit a high probability of a single shared causal variant. Concordant with previous studies (Pierce et al. 2018; Banovich et al. 2014), 56.1% of these eQTL-methylQTL pairs show an opposite effect direction. This proportion is 61.9% when only including pairs that had the same top-SNP based on QTL *p*-values, and 69.1% when further limiting to CpGs that are located on promoter regions. Pairs that show an opposite effect also show a high correlation between the absolute effect sizes (Pearson’s correlation 0.49, *p* < 2.2×10^-16^), while pairs with same effect directions don’t (Pearson’s correlation 0.03, *p*=0.64) (Figure S2). Colocalized CpGs located on promoters are more likely to show an opposite direction in effect with the gene than CpGs located outside promoters or enhancers (Fisher’s test *p*=3.835×10^-6^), but the same is not observed for CpGs located on enhancers when compared to those located outside promoters or enhancers (Fisher’s test *p*=0.6808).

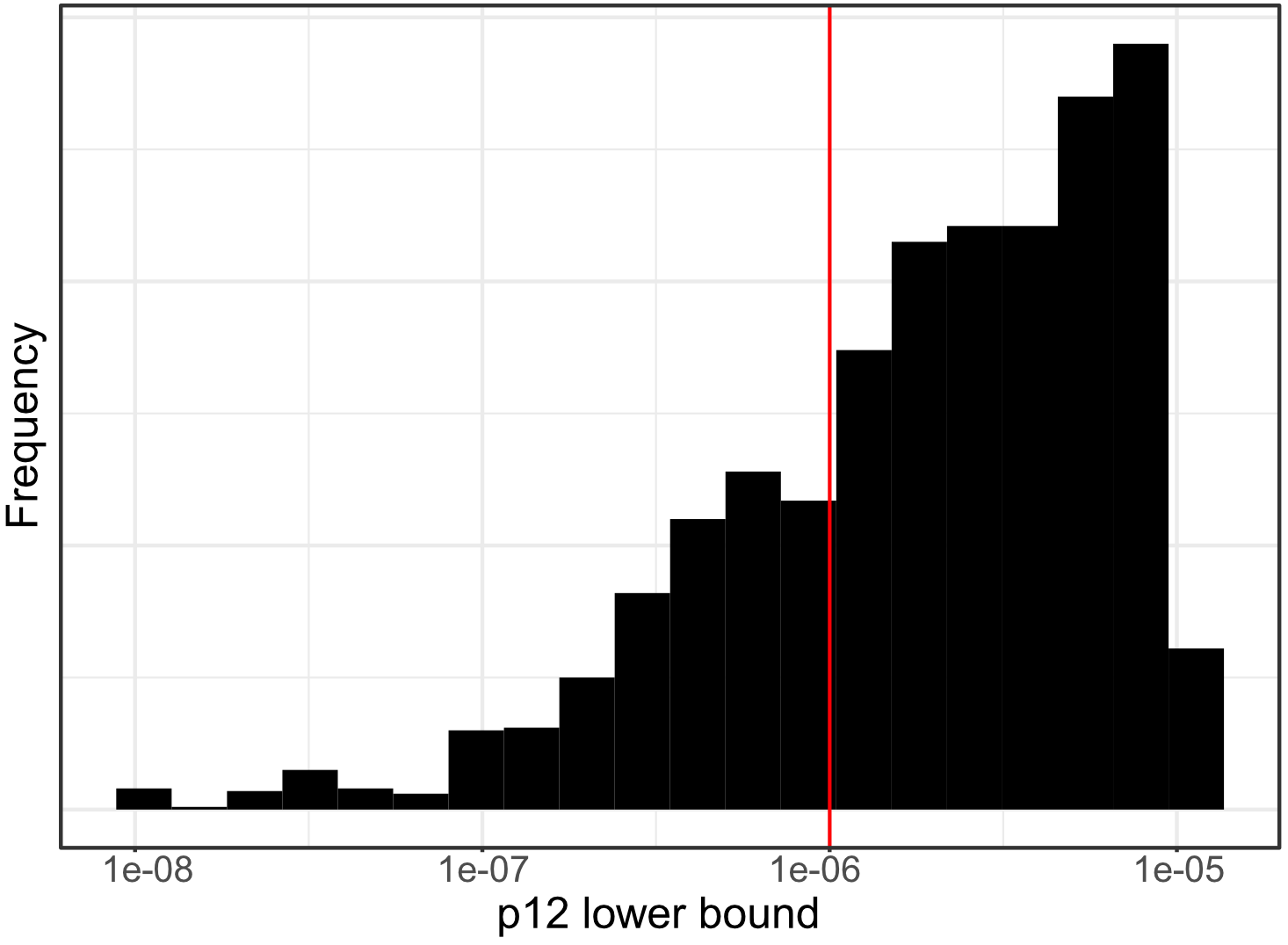

**Supplementary Figure S1.** Distribution of the lower bounds of the prior probabilities (p12) that suggest colocalization across 4,639 tested methylQTL-eQTL pairs. As the posterior probability for colocalization is dependent on the prior probability, a post-hoc sensitivity analysis was used to determine the range of prior probabilities for which colocalization is supported. Pairs passing the colocalization threshold with a range of p12 values from <1.0×10^−6^ to 1.0×10^−4^ (lower bound of p12 below 1.0×10^−6^) were considered to show robust support for colocalization.

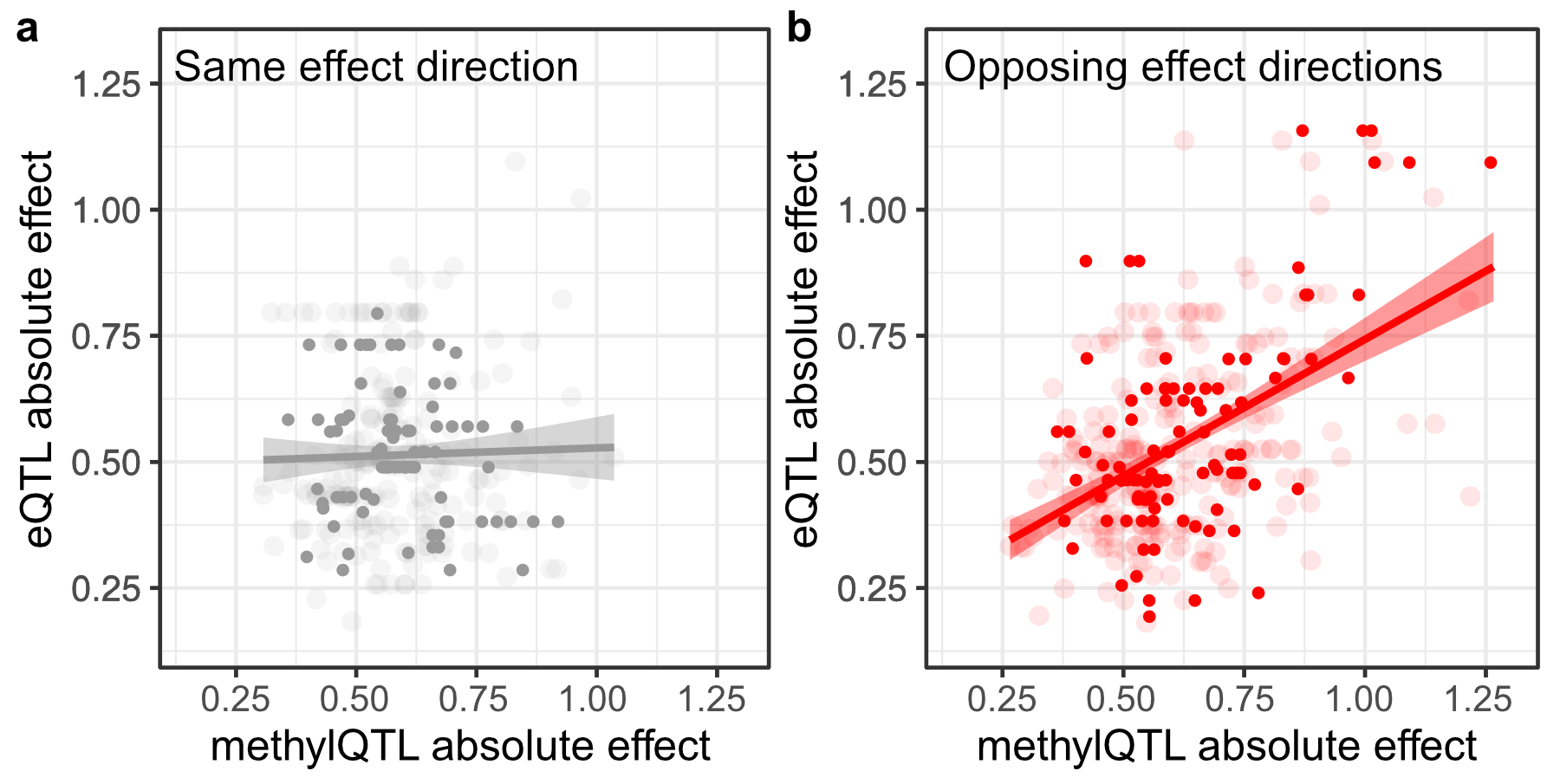

**Supplementary Figure S2. Genetically regulated promoter methylation alters target gene expression levels.** Relationship between the absolute effect sizes of colocalized methylQTLs and eQTLs that show the same direction of effect (**a**) and opposing direction of effect (**b**) on the target trait. Pairs that share the same top-SNP are plotted and variants located on promoter regions are highlighted. Smoothed means based on linear models in the form y ~ x and 95% confidence intervals are shown for each set.

**Supplementary Note 2. *Sharing of eQTLs between Indonesia and Europe.*** We compared the effect sizes of the colocalized and non-colocalized genes and the top-SNPs from each population. While the colocalized eQTLs show relatively large effect sizes for both population-specific top-SNPs in both populations, non-colocalized genes show overall lower effect sizes, and larger effect sizes for the population-specific top-SNP in each population. The eQTLs that were not colocalized and not significant eQTLs in the European population show overall lower effect sizes in the Indonesian data, compared to other tested eQTLs (mean

Indonesia-specific 0.32, mean other eQTLs 0.41, t-test *p*=9.4×10^-12^). The difference is even larger when comparing to the robust colocalized eQTLs (Figure 3a, means 0.32 and 0.52, t-test *p* < 2.2×10^-16^). This is concordant with the observation that eQTLs with larger effect sizes are more likely to be shared between populations (Stranger et al. 2012). Even though the larger sample size should allow the detection of lower effect eQTLs in the European data, we identify a subset of 166 Indonesia-specific eQTLs with relatively low effect sizes in our data. These genes did not exhibit any evidence of colocalization and did not have nominally significant eQTLs in the European data, indicative of true population-specific effects.

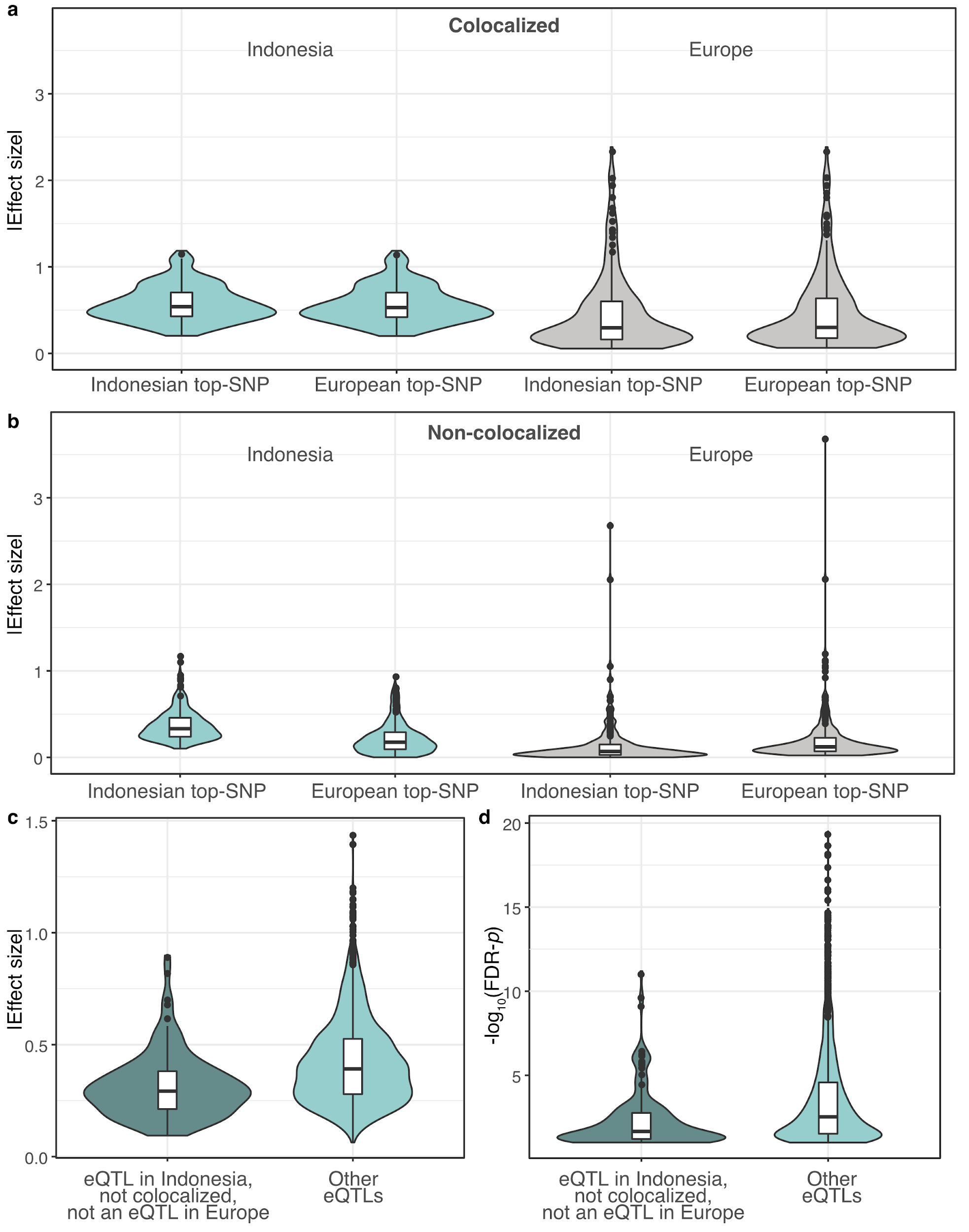

**Supplementary Figure S3.** Distributions of effect sizes of the colocalized (**a**) and non-colocalized (**b**) eQTLs where the same top-SNPs were tested in the Indonesian and European datasets. The effects of the top-SNPs in each population are shown. Indonesian effect sizes (**c**) and -log_10_(*p*-values) (**d**) of the eQTLs that don’t show any evidence of colocalization and don’t have significant SNPs in *cis* in the European data.

**Supplementary Note 3.** We compared the absolute effect sizes of the archaic ancestry driven QTLs and the effect sizes of the significant QTLs not driven by archaic ancestry. Denisovan and Neanderthal ancestry driven eQTLs (Supplementary Figure S4) and methylQTLs (Supplementary Figure S5) exhibit significantly larger absolute effect sizes than methylQTLs not driven by archaic ancestry. However, as the minor allele frequencies of the archaic driven QTLs are lower, we are less powered to detect small effect QTLs driven by archaic ancestry (Supplementary Figure S6). We performed allele frequency matching with the nearest neighbor matching method of the R package *MatchIt* v3.0.2 (Ho et al. 2011). There were no significant differences in the mean absolute effect sizes of the MAF matched sets and the archaic driven QTLs (Supplementary Figures S4 and S5).

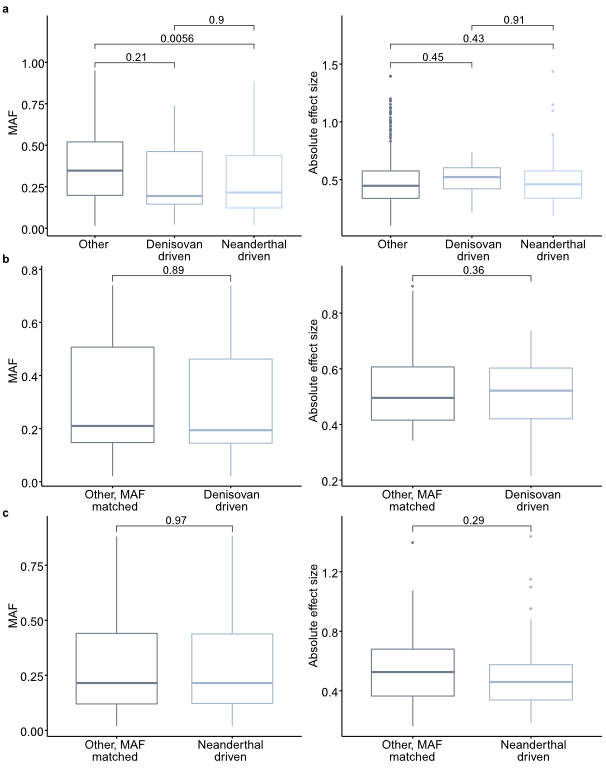

**Supplementary Figure S4.** Minor allele frequencies (MAF) and absolute effect sizes of eQTLs driven by Denisovan or Neanderthal introgression and eQTLs not driven by archaic introgression (“other”) before (**a**) and after (**b**, **c**) allele frequency matching. T-test p-values are indicated for each pairwise comparison.

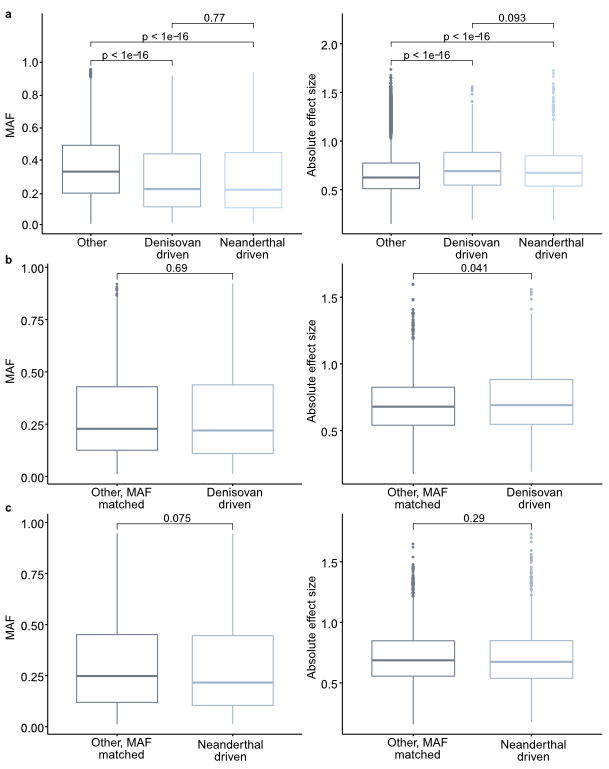

**Supplementary Figure S5.** Minor allele frequencies (MAF) and absolute effect sizes of methylQTLs driven by Denisovan or Neanderthal introgression and methylQTLs not driven by archaic introgression (“other”) before (**a**) and after (**b**, **c**) allele frequency matching. T-test p-values are indicated for each pairwise comparison.

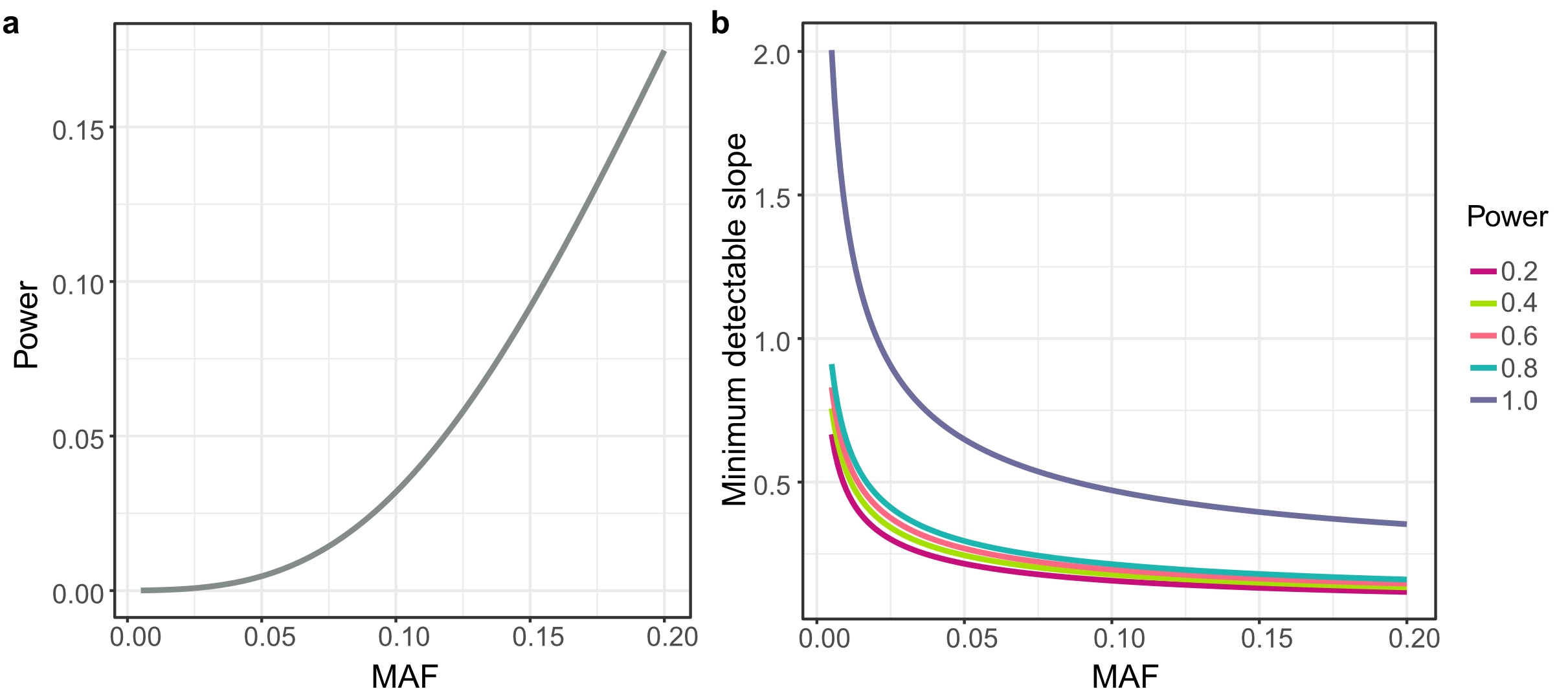

**Supplementary Figure S6.** **a:** Power to detect QTLs as a function of MAF when N=115. **b:** Minimum detectable slope in simple linear regression as a function of MAF, with various power levels. In both models, type I error rate was set to 0.01 and SD of the linear model to 0.2.

**Supplementary Note 4.** We assessed the credibility of the four Denisovan driven methylQTLs that colocalize with platelet count GWAS loci. First, we assessed our ability to correctly call genotypes on these positions, to correctly call methylQTLs, and to identify the correlation between the genotypes and the numbers of inferred Denisovan alleles. We used mappability scores generated with Umap (Karimzadeh et al. 2018) to assess mappability on regions overlapping these methylVariants. Umap calculates the single-read mappability of genome for a range of sequencing read lengths, the single-read mappability of a genomic region being defined as a fraction of that region which overlaps with at least one uniquely mappable k-mer. For a given sequence, mappability of 1 means that the sequence is uniquely mappable on the forward strand. Uniquely mappable regions with various kmers were downloaded from <https://bismap.hoffmanlab.org/>. All four variants are located on regions that are uniquely mappable with kmer lengths of 24, 36, 50, and 100bp, apart from chr6:29,799,383 which is on a region that is only uniquely mappable with kmers 36, 50 and 100bp. All four variants were called with high read depth, ranging from 29,492 to 37,373. All four variants have adequate MAFs, ranging from 0.161 to 0.302. All four methylQTLs show large effect sizes, the absolute effect size ranging from 0.66 to 0.88. Furthermore, all methylVariants show a clear correlation with the number of inferred Denisovan alleles, *R^2^* ranging from 0.73 to 0.90. The methylVariants associated with cg03118604 and cg03861427 are located within 741bp of each other and are in LD.

Then, we assessed whether sequence similarity across the genome could lead to spurious signals in the CpG methylation measurements using the Illumina EPIC array. We used megablast of BLASTN (Zhang et al. 2000) to map the forward sequences flanking the CpGs to the human reference genome. All four sequences map to the HLA locus with high confidence and do not map to other regions (Supplementary Table S10).
